# Impact of bacterial translocation in stroke outcome. Soluble CD14 as early clinical marker and effect of TLR4

**DOI:** 10.1101/2024.12.30.630676

**Authors:** Cristina Granados-Martinez, Nuria Alfageme-Lopez, Blanca Diaz-Benito, Ana Moraga, David Sevillano-Fernandez, Luis Alou-Cervera, M. Encarnación Fernandez-Valle, David Castejon-Ferrer, Patricia Calleja-Castaño, M. Angeles Moro, Macarena Hernandez-Jimenez, Ignacio Lizasoain, Jesus M. Pradillo

**Affiliations:** Pharmacology and Toxicology Department, School of Medicine, University Complutense of Madrid, Spain; Research Institute Hospital Universitario 12 de Octubre of Madrid, Spain; Cellular Biology Department, School of Medicine, University Complutense of Madrid, Spain; Microbiology Section, School of Medicine, University Complutense of Madrid, Spain; ICTS Complutense Bioimaging (BioImac), Research Assistance Centre, Complutense University of Madrid, Madrid, Spain; Neurovascular Pathophysiology, Cardiovascular Risk Factor and Brain Health Program, Centro Nacional de Investigaciones Cardiovasculares (CNIC), Madrid, Spain; AptaTargets S.L., Madrid, Spain

## Abstract

Bacterial infections are among the most common complications in stroke patients. While some factors triggering these infections are well established, bacterial translocation (BT) from the intestine to other organs represents another significant factor to consider. In our study, we observed a high percentage of animals with intestinal barrier dysfunction (GBD) and BT following stroke, with typical intestinal bacteria even detected in the lungs. Moreover, this process not only exacerbates peripheral/central inflammation but also increases lesion size. In this context, our data in stroke patients demonstrate the presence of GBD, associated with elevated levels of soluble CD14 as a marker, and its strong correlation with neurological status, infarct volume, and the development of infections. Finally, our findings highlight the neuroprotective effects of the absence or pharmacological inhibition of TLR4 using ApTOLL, which not only reduces infarct volume and inflammation but also mitigates GBD/BT processes following experimental stroke.

## Introduction

Stroke is one of the most devastating pathologies in terms of mortality, dementia, major adult disability and socioeconomic burden worldwide. In the European Union, stroke affects approximately 1.1 million individuals per year, making it the second leading cause of death and a major cause of adult disability^1^. Within its classification, two main types of stroke can be distinguished: ischemic stroke (IS) and hemorrhagic stroke. IS, which occurs more frequently, is treated primarily with clot fibrinolysis using tissue plasminogen activator (tPA) and/or mechanical clot removal via endovascular thrombectomy, therapies available to only a small percentage of patients^2^. In terms of stroke pathophysiology, during the first hours/days following the onset of IS, period known as the acute phase, inflammatory and immune responses play a key role in brain damage. These responses are triggered by the activation of immune receptors such as the Toll-like receptor family (TLRs) upon the release of damage-associated molecular patterns (DAMPs) from the ischemic brain. In this context, our group has demonstrated the crucial role of TLR4, a member of this receptor family, in stroke-related inflammation and ischemic damage. In fact, TLR4 knockout models have shown neuroprotection after experimental IS^3^. Moreover, we have investigated its role in hemorrhagic transformation following delayed tPA administration, as well as in neutrophil phenotype alterations^4, 5^. Additionally, we contributed to the development of a specific TLR4 antagonist, ApTOLL, which has shown potential to reduce inflammation and brain damage in experimental models^6^. This antagonist has now demonstrated both safety and efficacy in a recent Phase Ib/IIa clinical trial involving stroke patients^7^.

Once the acute phase is over, systemic immunosuppression (SI) may develop at both experimental and clinical levels^8, 9^. This SI is primarily due to the massive release of catecholamines, corticosteroids, and anti-inflammatory cytokines, resulting from the overactivation of the sympathetic nervous system (SNS) and the hypothalamic-pituitary-adrenal (HPA) axis^10–12^. These failures in the crosstalk between the damaged brain and the immune system facilitate complications that impair recovery and increase mortality. Among these, stroke-associated infections (SAIs) are particularly noteworthy, affecting 30-60% of patients, with urinary and respiratory tract infections being the most common^13–15^. Although several factors that promote the onset of these infections, such as SI, patient dysfunctions and hospital management are well established^16, 17^, their precise origin remains unclear. In recent years, however, both experimental and clinical evidence has pointed to alterations in the gastrointestinal (GI) microbiota, gut barrier damage (GBD), and the subsequent translocation of gut bacteria (BT) to organs like the lungs as a possible contributing factor to SAIs^18–21^. Furthermore, surrogate markers of GBD and BT, such as soluble CD14 (sCD14) and zonulin, have been identified in patients with various neurological conditions, including autism, stress and depression^22–24^. In stroke, previous studies have shown that sCD14, along with lipopolysaccharide-binding protein (LBP), can moderately predict the development of SAP in the clinic^25^. Specifically, CD14, which is normally expressed in a membrane-bound form on various monocyte subsets, is released in its soluble form after enzymatic shedding upon recognition of pathogen-associated molecular patterns (PAMPs), such as lipopolysaccharides (LPS), due to increased gut permeability and/or BT^24, 26–28^. Thus, detecting these markers of GBD/BT in stroke patients could provide valuable insight into the early prediction of SAIs with a gut origin, potentially enabling more effective therapeutic interventions to improve patient outcomes.

Finally, imaging techniques like MRI play a central role in the clinical diagnosis of stroke^29^. MRI offers unique opportunities for obtaining non-invasive structural and functional information about various organs, including the intestine, and allows the use of contrast agents (CAs) to enhance the visualization of different structures. While few studies have focused on analyzing GBD/BT using abdominal MRI in the context of stroke, certain contrast agents, such as Manganese Chloride (MnCl2), which is paramagnetic and exclusively excreted via the hepatobiliary system after oral administration, may be particularly useful for early prediction of these post-stroke complications^30^.

With this background, our aim was to investigate the impact of BT on the inflammatory response and ischemic damage experimentally, to develop a novel abdominal MRI protocol for assessing GBD after stroke, and to identify plasma markers of GBD/BT for the early detection of these processes in both clinical and experimental settings. Additionally, we sought to analyze the effect of lesion size on GBD/BT development, and, more interestingly, the impact of TLR4 deficiency or pharmacological inhibition with ApTOLL on these processes following experimental stroke.

## Results

### IS induces GBD and BT

To investigate BT in the IS model (MCAO), we analyzed intestinal bacterial growth by culturing samples from peritoneal organs and lungs of *naïve*, and at 72h post-surgery of sham, and MCAO rats. No bacterial growth was detected in *naïve* or sham groups (Supp. Table1a), but 75% of MCAO animals showed bacterial growth mainly in the mesenteric lymph nodes (MLN; Supp. Table1b), and in liver and spleen, and 76% of MCAO-BT rats exhibited bacterial growth in the lungs (Fig. 1a). Gram-positive bacteria were more common than Gram-negative bacteria, with Bacillota and *Actinomycetota* being the most abundant Gram-positive phyla, while *Pseudomonadota* and *Bacteroidota* predominated among Gram-negative bacteria (Fig. 1b, c). The bacteria in the lungs of MCAO-BT animals included *Lactobacillus sp.*, *Enterococcus sp.*, *Clostridium sp.*, and *Escherichia coli*, which are associated with intestinal microbiota and linked to respiratory infections in stroke patients (Fig. 1d). Using contrast-enhanced MRI, we observed significantly increased GBD by detecting CAs in the spleen and MLN of MCAO-BT animals compared to MCAO-NBT and *naïve* groups (Fig. 1e). Immunofluorescence analysis of the gut showed that MCAO-BT animals had reduced ZO-1 levels (tight junction protein) and increased MMP-9 expression (inflammation/barrier disruption) compared to controls (Fig. 1f, g). These findings highlight that IS induces GBD, leading to BT from the intestine to peritoneal organs and lungs in a significant proportion of animals.

**Figure 1.**
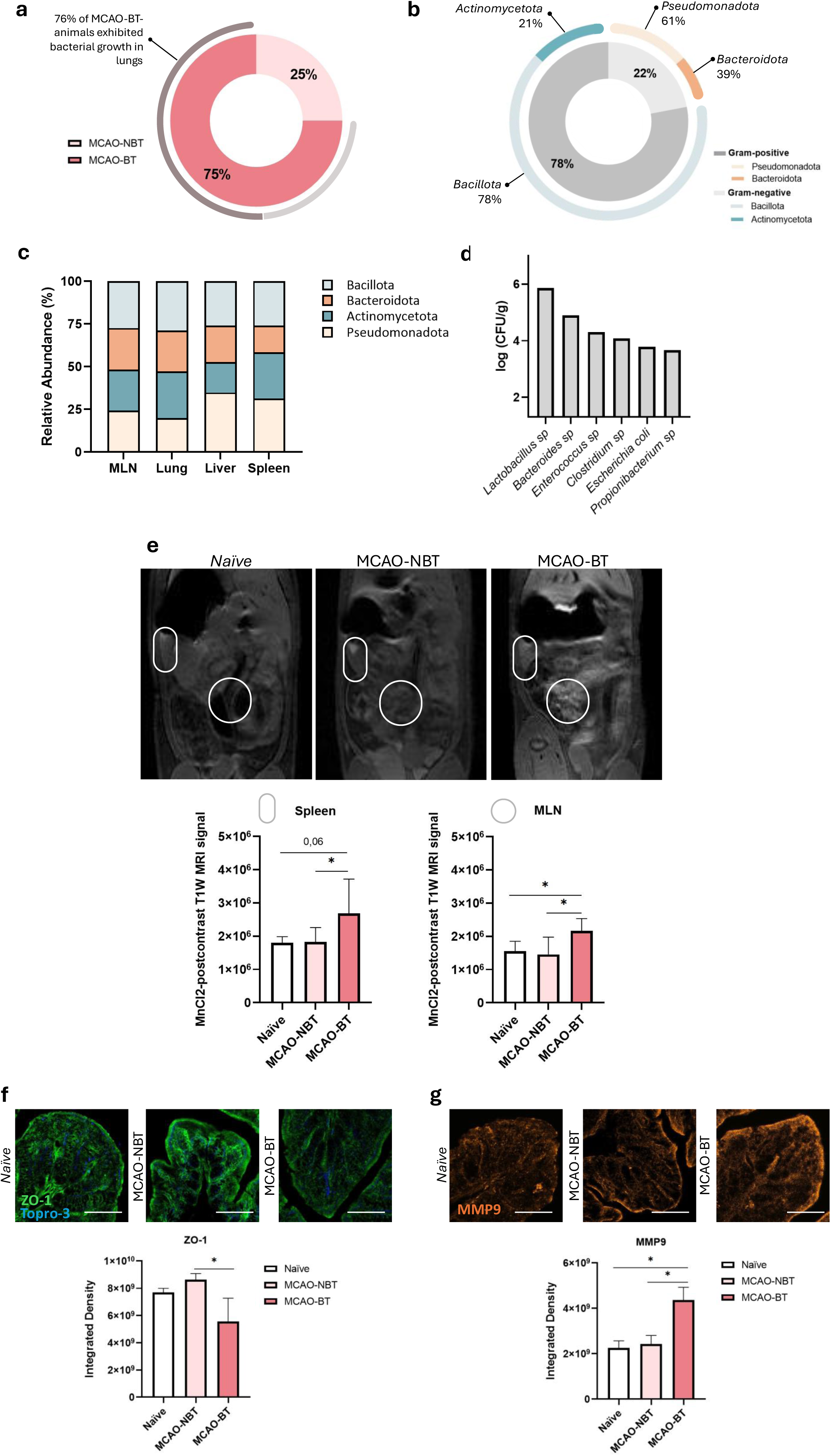
Bacterial translocation (BT) and gut barrier disruption (GBD) following experimental ischemic stroke. **(a)** Percentage of IS animals with (MCAO-BT, n=21) or without (MCAO-NBT, n=7) BT at 72h post-surgery. Bacterial growth in the lungs of MCAO-BT group (n=16) is also shown. **(b)** Relative abundance of Gram-positive and Gram-negative bacteria, and bacterial phyla isolated from microbiological cultures of organs at 72h after experimental IS. **(c)** Relative abundance of bacterial phyla isolated from microbiological cultures of MLN, lung, liver, and spleen. **(d)** Representation of the log CFU/g of bacterial species isolated from the lung. **(e)** Intestinal permeability at 72h post-ischemic stroke assessed by MRI with contrast agent in *naïve* (n=7), MCAO-NBT (n=7), and MCAO-BT (n=21) groups. **(e, top)** Representative abdominal T1W-MRI images showing the spleen (oval) and MLN (circle)**. (e, bottom)** MnCl2-postcontrast T1W-MRI signal in spleen and MLN. Results are shown as mean ± SD; one-way ANOVA followed by Tukey’s post-hoc test, \**p* < 0.05. **(f and g)** Immunofluorescence analysis of GBD in naïve (n=4) and at 72h after IS in MCAO-NBT (n=6), and MCAO-BT (n=6) animals. **(f)** Confocal images of **(f)** ZO-1 (green) and **(g)** MMP9 (red) expression. Scale bar: 50 μm. Representation of the integrated density of ZO-1 and MMP9 signals is shown. Results are shown as mean ± SD; one-way ANOVA followed by Tukey’s post-hoc test, \**p*<0.05.

### Effect of GBD/BT on peripheral, central inflammation and infarct volume

The effect of GBD/BT effect on peripheral inflammation, assessed by flow cytometry, showed a significant reduction in B cells in the bone marrow (BM) and a trend toward reduced monocytes and increased granulocytes in MCAO-BT animals compared to MCAO-NBT and *naïve* groups (Fig. 2a). In peripheral blood, BT after IS led to reduced T-CD4+ and T-CD8+ lymphocytes, increased monocyte levels, and a trend toward reduced granulocytes, with significant lymphopenia observed only in the MCAO-BT group (Fig. 2b). No differences were noted in the MNL, spleen, or lung cell populations (data not shown). In plasma, GBD/BT following IS increased the proinflammatory chemokines CINC-1 and CCL20, as well as the anti-inflammatory cytokines IL-1Ra and IL-10, compared primarily to the *naïve* group (Fig. 2c). Central inflammation analysis showed IS effects at 72h post-surgery in microglial activation and granulocyte infiltration. Additionally, CD3+ lymphocyte infiltration was significantly higher in MCAO-BT animals compared to the MCAO-NBT group (Fig. 2d-f). Finally, infarct volume at 72h was significantly higher in the MCAO-BT group compared to MCAO-NBT, indicating that BT worsens both peripheral/central inflammation and exacerbates ischemic injury (Fig. 2g). Among MCAO-BT animals, two subgroups emerged based on infarct size—above or below the mean—with infarct size appearing to influence the likelihood of BT development (Fig. 2h).

**Figure 2.**
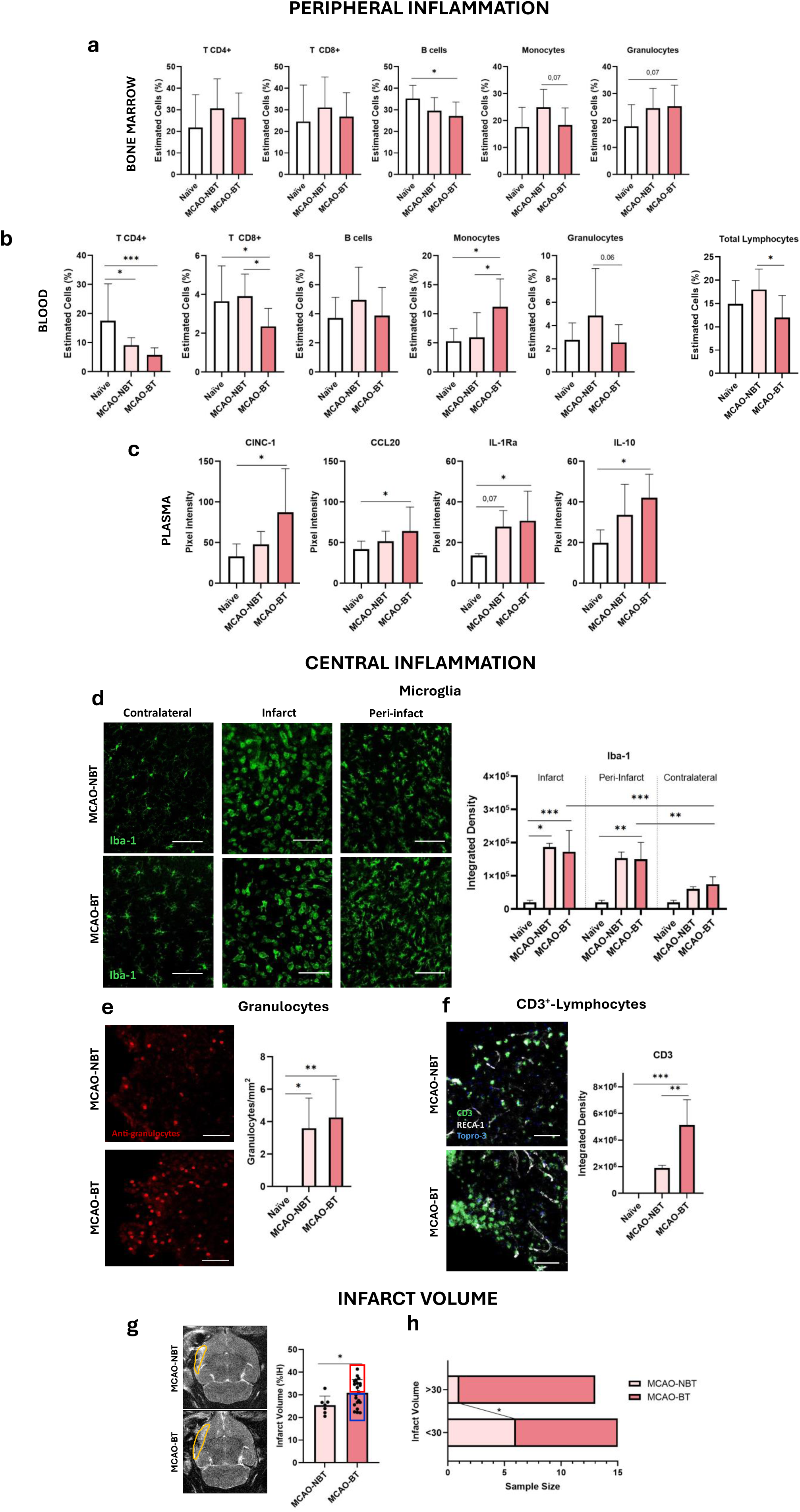
Effect of GBD/BT on peripheral, central inflammation and infarct volume. **(a-c)** Immune cell levels assessed by flow cytometry in **(a)** bone marrow and **(b-c)** peripheral blood in *naïve* (n=8) and at 72h post-surgery in MCAO-NBT (n=7), and MCAO-BT (n=21) animals. Results are shown as mean ± SD; one-way ANOVA followed by Tukey’s post-hoc test, \**p*<0.05, \*\*\**p*<0.001. **(c)** Plasma levels analyzed by cytokine/chemokine array of CINC-1, CCL20, IL-1ra, and IL-10 in *naïve* (n=3) and at 72h post-surgery in MCAO-NBT (n=6-8), and MCAO-BT (n=6-8) groups. Results are shown as mean ± SD; Kruskal-Wallis test followed by Dunn’s post-hoc test, \**p*<0.05. **(d)** Microglial activation in ischemic brain tissue. **(d, top)** Representative confocal images of Iba-1 (green) expression in *naïve* (n=4) and MCAO-NBT (n=7), and MCAO-BT (n=21) groups. Scale bar: 50 μm. **(d, bottom)** Quantification of Iba-1 levels in infarct, peri-infarct, and contralateral hemispheres. Results are shown as mean ± SD; Kruskal-Wallis test followed by Dunn’s post-hoc test, \**p*<0.05, \**p*<0.01, \*\**p*<0.001. **(e)** Granulocyte infiltration in the ischemic brain. **(e, left)** Representative confocal images showing granulocyte (red) infiltration in ischemic brain parenchyma at 72h post-surgery in MCAO-NBT (n=7) and MCAO-BT (n=21) groups. Scale bar: 50 μm. **(e, right)** Quantification of granulocytes per mm^2^ in the ipsilateral hemisphere. Results are shown as mean ± SD; Student’s t-test, \**p*<0.05, \*\**p*<0.01. **(f)** CD3+-lymphocyte infiltration in the ischemic brain. **(f, left)** Representative confocal images of CD3 (green), RECA-1 (white), and Topro-3 (blue) expression in ischemic brain tissue at 72h post-surgery in MCAO-NBT (n=6) and MCAO-BT (n=6) groups. Scale bar: 50 μm. **(f, right)** Quantification of CD3+ T-cells in the infarcted region. Results are shown as mean ± SD; Student’s t-test, \*\**p*<0.01, \*\*\**p*<0.001. **(g)** Infarct volume at 72h post-surgery in MCAO-NBT (n=7) and MCAO-BT (n=21) animals. Infarct volume is expressed as the percentage of the ipsilateral hemisphere (%IH). Results are shown as mean ± SD; Student’s t-test, *p* < 0.05. Infarct volumes > 30% (red) and < 30% (blue) are indicated. **(h)** Proportion of ischemic animals with infarct volumes greater or less than 30% of the IH and whether BT occurred. Chi-square test, \**p*<0.05. Infarct volume > 30%: n (MCAO-NBT)=1, n (MCAO-BT)=12; Infarct volume < 30%: n (MCAO-NBT)=6, n (MCAO-BT)=9.

### sCD14 as early clinical/experimental marker of GBD/BT and SAIs

To validate our experimental findings on GBD/BT in a clinical setting, we measured circulating levels of zonulin and sCD14 by ELISA as markers of GBD 24h after admission in a cohort of 152 ischemic stroke patients (baseline characteristics and patient grouping based on sCD14 levels are provided in Supplemental Table 2). Although our results for zonulin were inconclusive (data not shown), the findings for sCD14 were positive. After stratifying the patients into tertiles based on sCD14 levels, the means and probabilities of various clinical variables were calculated, adjusting for potential confounders such as gender, age, and dyslipidemia. Our results showed a significant association between higher sCD14 levels and larger infarct volumes (Fig. 3a and Suppl Table 3). Patients with the highest sCD14 levels also had impaired collateral circulation, worse clinical outcomes, determined by NIHSS score at admission and at 24h (Fig. 3b,c and Suppl Table 3). Finally, the probability of in-hospital infections stratified by sCD14 levels showed a significant increase in the group with the highest levels of this marker, compared to the other groups (Fig. 3d and Suppl Table 3). These findings suggest that elevated sCD14 levels are strongly associated with worse stroke outcomes and a higher risk of SAIs. Additionally, sCD14 might serve as an early biomarker for identifying patients at risk of GBD, emphasizing its potential clinical utility.

**Figure 3.**
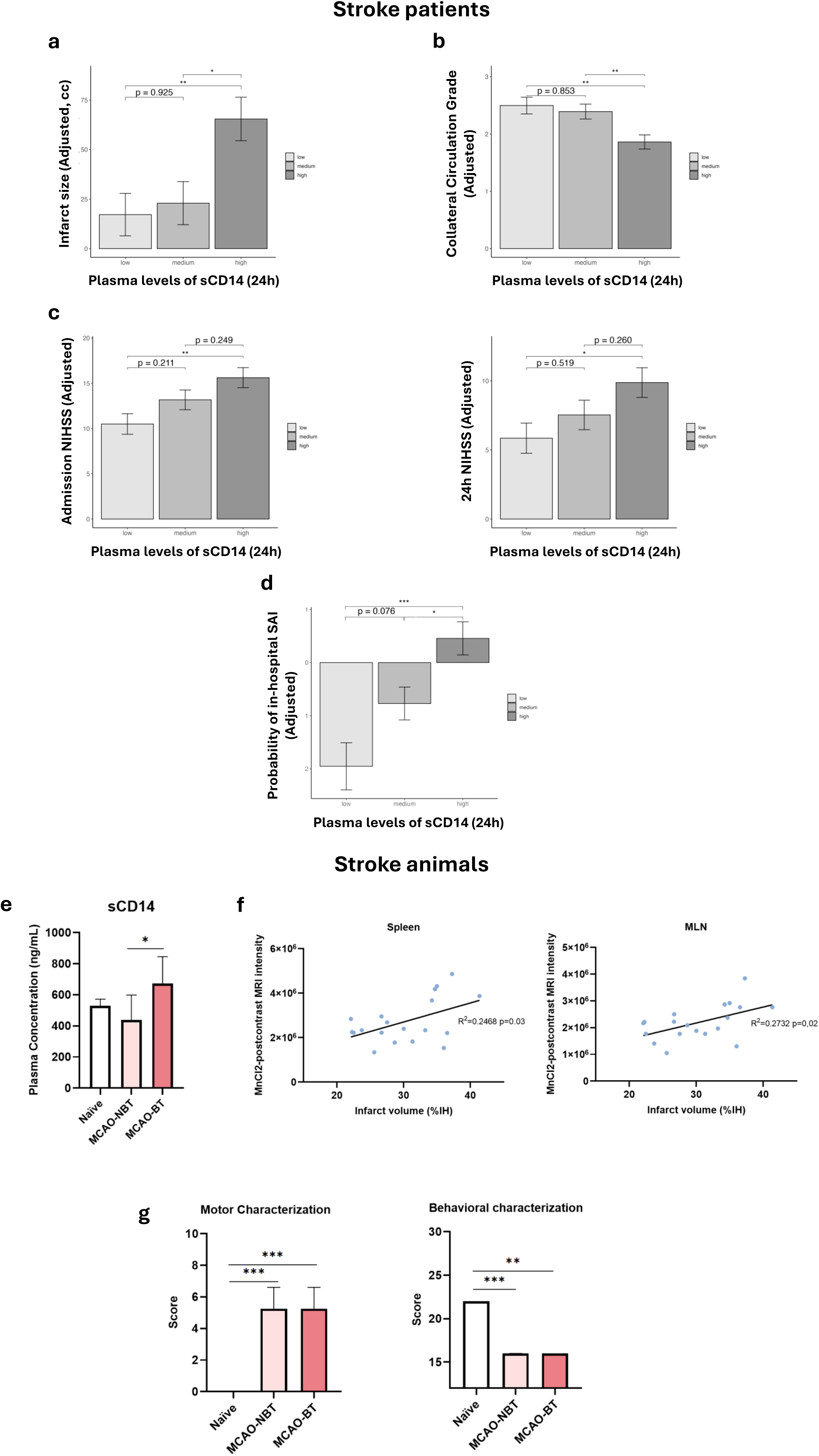
sCD14 as early clinical and experimental marker of GBD, BT and SAIs. **(a–d)** Adjusted mean values and probabilities for clinical variables, stratified by CD14 levels at 24h of admission (low, medium, high). Analyses include **(a)** infarct volume, **(b)** collateral circulation status, **(c)** baseline NIHSS and NIHSS at 24h, and **(d)** probability of hospital-acquired infections. Bars represent adjusted mean values or probabilities derived from multivariable models, with error bars indicating 95% confidence intervals. Pairwise comparisons between CD14 levels are indicated with *p* values above corresponding groups. Adjustments were made for covariates including sex, age, and dyslipidemia to explore associations between CD14 levels and key clinical outcomes. **(e)** Plasma levels of sCD14 (ng/mL) in *naïve* group (n=5) and at 72 h in the MCAO-NBT and MCAO-BT groups (n=12). Results are shown as mean ± SD; one-way ANOVA followed by Tukey’s multiple comparisons test, \**p*<0.05. **(f)** Correlation between infarct volume and MnCl2-postcontrast MRI intensity observed in **(f, left)** spleen (R^2^ = 0.2468, *p* = 0.03) and **(f, right)** MLN (R^2^ = 0.2732, *p* = 0.02). **(g)** Effect of stroke and BT on neurological deficits assessed in naïve animals (n=6) and at 72h post-surgery in MCAO-NBT and MCAO-BT groups (n=11). **(g, left)** Motor function scale. **(g, right)** Behavioral scale. Results are expressed as mean ± SD; Kruskal-Wallis test followed by Dunn’s multiple comparisons test, \*\*\**p<0.001*.

In our animal model, the levels of sCD14 analyzed in *naïve* and in MCAO-NBT and BT groups, revealed a significant increase in this marker in IS animals with BT compared to the other experimental groups (Fig. 3e). To better understand the relationship between infarct volume, sCD14 levels and GBD/BT, the correlation between infarct volume and CAs intensity in abdominal MRI-images, showed a positive correlation between these two variables in both the spleen and MLN within the MCAO-BT group (Fig. 3f). In contrast, analyzing the stroke outcome in our experimental model, only a worsening due to the stroke itself was observed (Fig. 3g). These results, consistent with patient data, confirm that ischemic animals with BT have larger infarct volumes and increased intestinal barrier permeability, as shown by elevated plasma sCD14 levels and CAs accumulation in peritoneal organs.

### Pharmacological TLR4 inhibition with ApTOLL reduces GBD/BT and improves stroke outcome

To confirm the neuroprotective effect of ApTOLL, infarct volume and neurological function were assessed in *naïve*/MCAO-Vehicle and *naïve*/MCAO-ApTOLL-treated groups at 72 hours post-surgery. The results demonstrated that pharmacological TLR4 inhibition improved stroke outcomes in the ApTOLL-treated group compared to the vehicle group (Fig. 4a; Suppl. Fig. 1a, b). When evaluating the effect of ApTOLL on BT in this cohort, we observed a non-significant reduction in BT in the MCAO-ApTOLL group compared to the MCAO-Vehicle group. Similarly, bacterial growth in the lungs was slightly reduced in the ApTOLL-treated group, though this difference was not statistically significant (Fig. 4b). Notably, ApTOLL treatment increased the percentage of Gram-positive bacteria and decreased Gram-negative bacteria compared to the vehicle group. In the MCAO-ApTOLL group, *Actinomycetota* predominated among Gram-positive bacteria, while only *Pseudomonadota* was present among Gram-negative bacteria. In contrast, the MCAO-Vehicle group contained both *Pseudomonadota* and *Bacteriodota* (Fig. 4c). When infarct volume was analyzed based on treatment group and BT status, a reduction in lesion size was observed only in MCAO-ApTOLL animals without BT, compared to the vehicle-treated groups (Fig. 4d). Intestinal barrier analysis revealed decreased ZO-1 expression in animals with BT, regardless of treatment (Fig. 4e). Additionally, gut-MMP9 levels were significantly higher in the MCAO-Vehicle-BT group compared to the *naïve*, MCAO-Vehicle-NBT, and MCAO-ApTOLL-BT groups (Fig. 4f).

**Figure 4.**
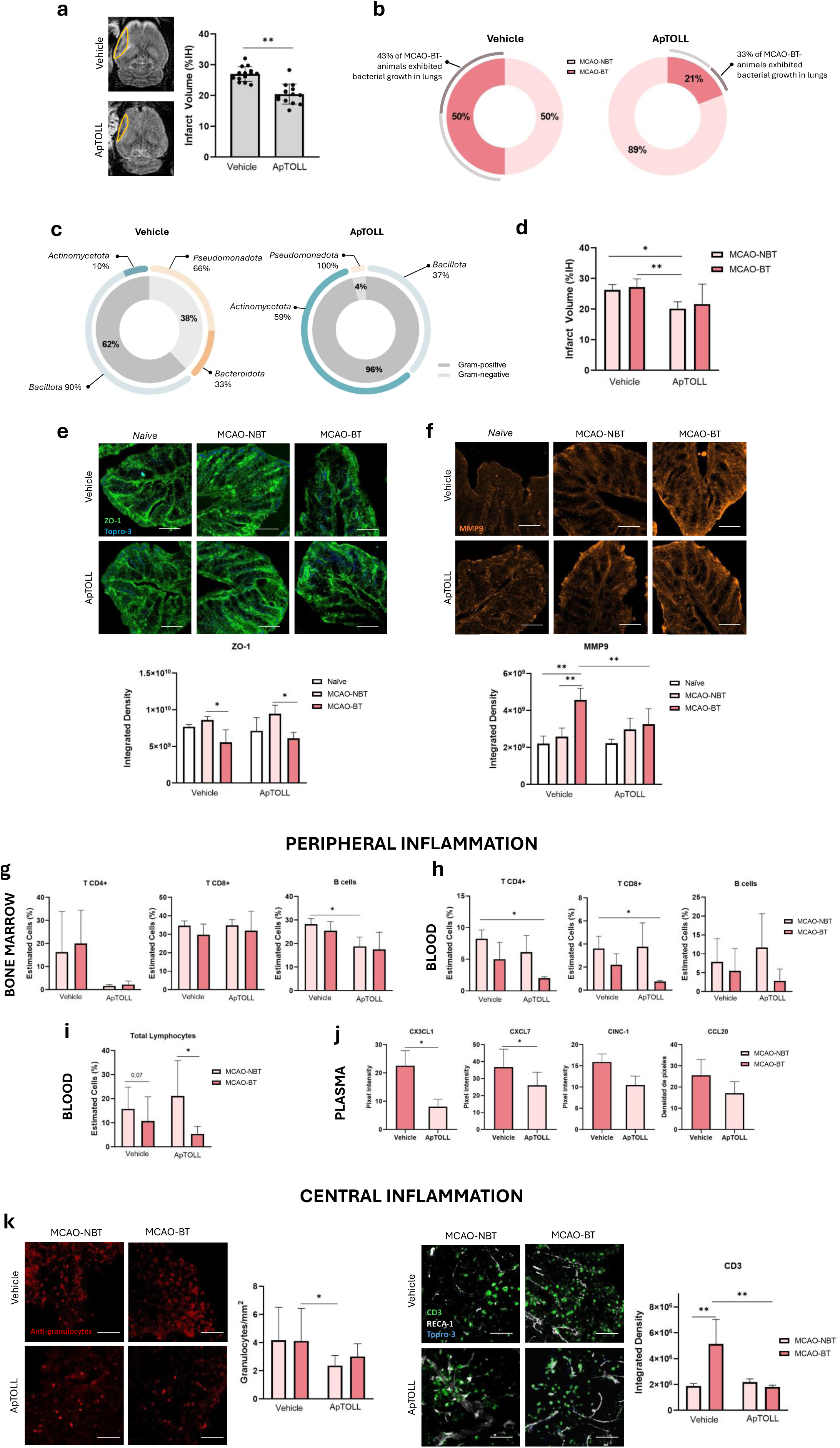
Effect of ApTOLL treatment on stroke outcome, BT, and immune responses. **(a)** Infarct volume at 72h post-surgery according to treatment (Vehicle, ApTOLL; n=14). Infarct volume is expressed as the percentage of the ipsilateral hemisphere (%IH). Results are shown as mean ± SD; Student’s t-test, \*\**p*<0.01. **(b)** Proportion of ischemic animals developing (MCAO-BT) or not (MCAO-NBT) BT in the Vehicle-treated group (n (MCAO-NBT)=7, n (MCAO-BT)=7) and ApTOLL-treated group (n (MCAO-NBT)=11, n (MCAO-BT)=3). **(c)** Relative abundance of gram-positive and gram-negative bacteria and bacterial phyla isolated from microbiological cultures of organs at 72h post-surgery in Vehicle and ApTOLL-treated groups. **(d)** Infarct volume at 72h post-surgery according to treatment and BT status (Vehicle: n (MCAO-NBT)=7, n (MCAO-BT)=7; ApTOLL: n (MCAO-NBT)=11, n (MCAO-BT)=3). Results are shown as mean ± SD; Kruskal-Wallis test followed by Dunn’s post-hoc test, \**p*<0.05, \*\**p*<0.01. **(e)** ZO-1 and **(f)** MMP9 expression in colon tissue. **(e, top)** Representative confocal images of ZO-1 (green) and (f, top) MMP9 expression (red). Scale bar: 50 μm. **(e, bottom)** Quantification of ZO-1 and **(f, bottom)** MMP9 signal expressed as integrated density in *naïve* Vehicle- and ApTOLL-treated animals (n=4) and ischemic Vehicle-treated (n (MCAO-NBT)=6, n (MCAO-BT)=6) and ApTOLL-treated (n (MCAO-NBT)=7, n (MCAO-BT)=3) groups. Results are shown as mean ± SD; Kruskal-Wallis test followed by Dunn’s post-hoc test, \**p*<0.05, \*\**p*<0.01. **(g-i)** Immune cell levels in **(g)** BM and **(h,i)** peripheral blood at 72h post-surgery by flow cytometry in ischemic animals treated with Vehicle (n (MCAO-NBT)=7, n (MCAO-BT)=7) or ApTOLL (n (MCAO-NBT)=11, n (MCAO-BT)=3). Results are shown as mean ± SD; Kruskal-Wallis test followed by Dunn’s post-hoc test, \**p*<0.05. **(j)** Plasma levels of CX3CL1, CXCL7, CINC-1, and CCL20 analyzed by cytokine/chemokine array in Vehicle MCAO-BT (n=3) and ApTOLL MCAO-NBT (n=7) animals. Results are shown as mean ± SD; Mann-Whitney test, \**p*<0.05. **(k)** Immune cell infiltration in ischemic brain at 72h post-surgery. **(k, left)** Granulocyte infiltration assessed by immunofluorescence in ischemic brain from Vehicle MCAO-NBT and MCAO-BT (n=6) and ApTOLL MCAO-NBT (n=5) and MCAO-BT (n=3) animals. Representative confocal images of granulocyte staining (red) in infarct region. Scale bar: 50 μm. Quantification of infiltrated granulocytes/mm² in the lesion area. Results are shown as mean ± SD; Kruskal-Wallis test followed by Dunn’s post-hoc test, \**p*<0.05. **(k, right)** T-cell infiltration assessed by immunofluorescence in ischemic brain from Vehicle MCAO-NBT and MCAO-BT (n=6) and ApTOLL MCAO-NBT (n=5) and MCAO-BT (n=3) animals. Representative confocal images of CD3 (green), RECA-1 (white), and Topro-3 (blue) staining in infarct region. Scale bar: 50 μm. Quantification of T-cell infiltration expressed as integrated density. Results are shown as mean ± SD; Kruskal-Wallis test followed by Dunn’s post-hoc test, \**p*<0.05, \*\**p*<0.01.

It is important to note that no significant differences in infarct volume were observed when comparing vehicle-treated animals to the previously untreated cohort (Supp. Fig. 1c). To further investigate ApTOLL’s effect on BT, we compared BT percentages between untreated animals, the MCAO-Vehicle group, and the MCAO-ApTOLL group. This analysis revealed a significant reduction in BT in the ApTOLL-treated group, along with a non-significant reduction in bacterial growth in the lungs compared to the other groups (Suppl. Fig. 2a, b). Furthermore, infarct volume and neurological function analyses showed a significant reduction in lesion size and improved stroke outcomes in the MCAO-ApTOLL group, regardless of BT status, compared to both the untreated and MCAO-Vehicle groups (Supp. Fig. 2c, d).

The evaluation of ApTOLL’s effect on immune cell populations in BM and blood revealed no immune alterations in the *naïve*-ApTOLL group compared to the vehicle group. However, in IS animals, ApTOLL treatment improved the immune response, particularly in T-CD4+ cells in BM and T-CD8+ cells, B cells, and monocytes in blood (Supp. Fig. 3a, b, c). In IS animals based on the BT process, ApTOLL treatment reduced B cells in BM and decreased T-CD4+ and T-CD8+ cells in peripheral blood in the MCAO-ApTOLL-BT group compared to the MCAO-Vehicle-NBT group (Fig. 4g, h). Both ApTOLL and vehicle BT groups showed a reduction in total lymphocyte percentages compared to their respective NBT groups (Fig. 4i). No differences were observed in immune populations in the MLN, spleen, or lungs (data not shown). Regarding chemokines/cytokines, plasma from the MCAO-Vehicle-BT and MCAO-ApTOLL-NBT groups revealed a significant reduction in proinflammatory chemokines CX3CL1 and CXCL7 in the ApTOLL-NBT group compared to the vehicle-BT group, with a non-significant reduction in CINC1 and CCL20 levels (Fig. 4j). Finally, ApTOLL’s effect on central inflammation in the context of stroke and BT showed a reduction in granulocyte and, notably, T-cell infiltration in the ischemic brain in the ApTOLL-treated groups compared to the MCAO-Vehicle-BT group (Fig. 4K).

In summary, these results demonstrate that pharmacological inhibition of TLR4 by ApTOLL after experimental stroke reduces infarct volume, BT, GBD, and both peripheral and central inflammation.

### Effect of TLR4 deficiency and infarct size on BT/GDB

We first assessed the impact of proximal versus distal MCAO and the effect of TLR4 deficiency on infarct volume. Our results revealed a significant reduction in lesion size in the WT-distal group compared to the WT-proximal group. Regarding TLR4 deficiency, infarct volume was significantly reduced in the WT and TLR4-KO groups with the same MCAO location (WT-proximal vs. TLR4-KO). However, when comparing lesion size between the WT-distal and TLR4-KO groups, the TLR4-deficient group exhibited a significant increase in lesion size (Fig. 5a).

**Figure 5.**
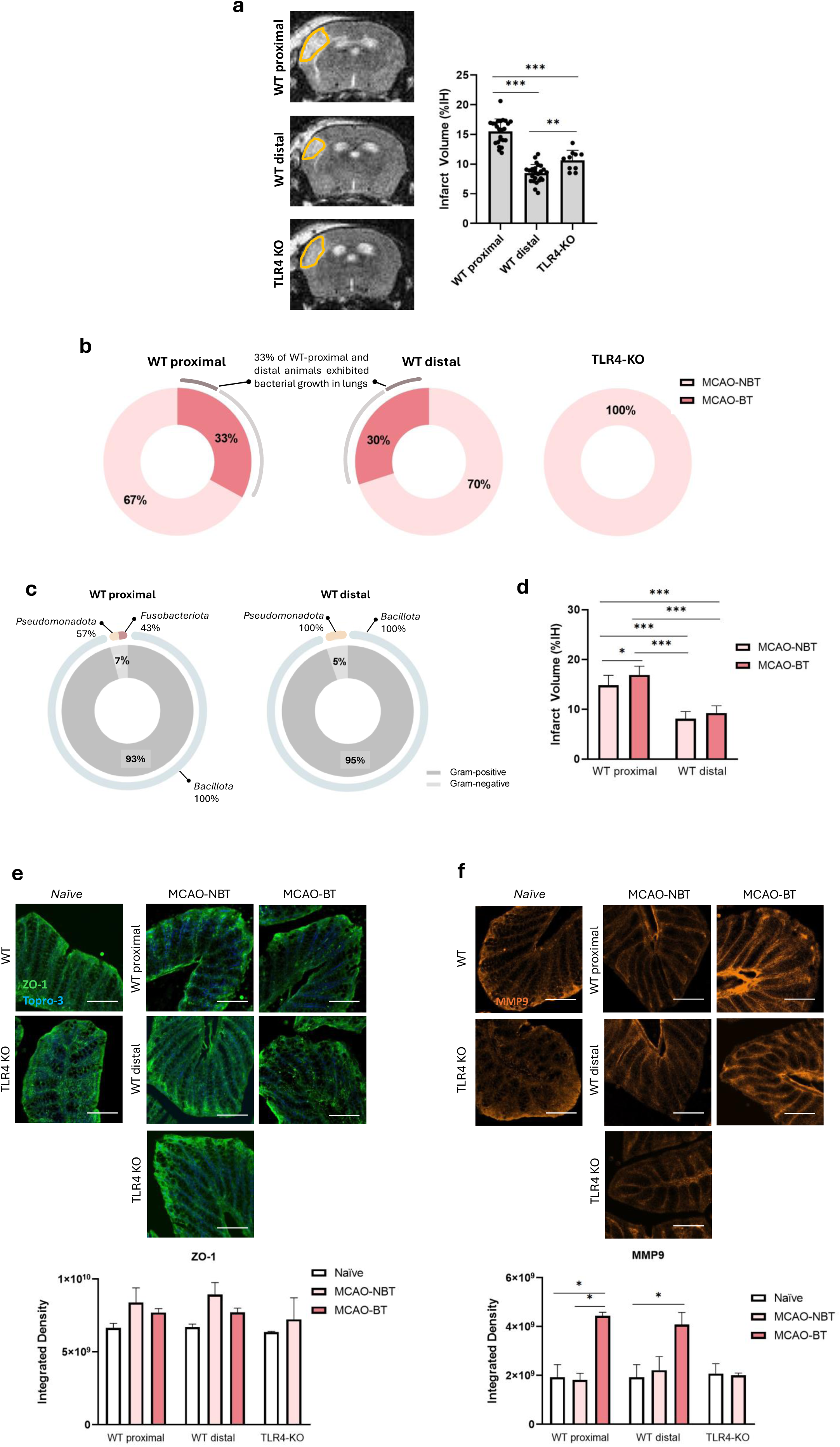
Role of TLR4 in infarct volume, BT, and intestinal barrier dysfunction following ischemic stroke. **(a)** Infarct volume at 72h post-surgery in WT proximal (n=26), WT distal (n=24), and TLR4-KO (n=10) groups. The infarcted area is shown in yellow. Infarct volume is expressed as the percentage of the ipsilateral hemisphere (%IH). Results are shown as mean ± SD; one-way ANOVA followed by Tukey’s multiple comparisons test, \*\**p*<0.01, \*\*\**p*<0.001. **(b)** Proportion of ischemic animals developing (MCAO-BT) or not (MCAO-NBT) BT at 72h post-surgery in WT proximal (n (MCAO-NBT)=13, n (MCAO-BT)=11), WT distal (n (MCAO-NBT)=16, n (MCAO-BT)=10), and TLR4-KO (n=10) groups. **(c)** Relative abundance of gram-positive and gram-negative bacteria and bacterial phyla isolated from microbiological cultures of organs at 72h post-surgery in WT proximal and WT distal groups. **(d)** Infarct volume at 72h post-surgery in WT proximal MCAO-NBT (n=18) and MCAO-BT (n=8), and WT distal MCAO-NBT (n=16) and MCAO-BT (n=8) mice. Results are shown as mean ± SD; two-way ANOVA followed by Bonferroni’s multiple comparisons test, \**p*<0.05, ***p<0,001. **(e)** ZO-1 and **(f)** MMP9 levels in colon tissue at 72h post-surgery assessed by immunofluorescence. **(e, top)** Representative confocal images of ZO-1 (green) and Topro-3 (blue), **(f,top)** and MMP9 (red) staining in colon from WT *naïve*, WT proximal MCAO-NBT and MCAO-BT, WT distal MCAO-NBT and MCAO-BT, TLR4-KO *naïve*, and TLR4-KO (n=6) groups. Scale bar: 50 μm. **(e, bottom)** Quantification of ZO-1 and MMP9 signal expressed as integrated density in the same groups. Results are shown as mean ± SD. Kruskal-Wallis test followed by Dunn’s multiple comparisons test, \**p*<0.05.

Analysis of the impact of infarct size on BT showed a non-significant reduction between the WT-proximal and distal groups, with no differences in bacterial growth in the lungs. However, TLR4 deficiency completely prevented BT development in the TLR4-KO group (Fig. 5b). Evaluation of bacterial types outside the intestine in the WT groups with BT revealed no significant differences in Gram-positive/negative bacteria or bacterial phyla between the WT-proximal and distal groups (Fig. 5c). Infarct volume analysis in WT groups based on BT showed an increase in lesion size in the WT-proximal group with BT compared to those without. Moreover, the WT-distal group exhibited smaller infarct volumes, regardless of BT status, compared to both WT-proximal groups (Fig. 5d).

Finally, intestinal barrier integrity (ZO-1 expression) showed no significant differences between *naïve* and IS groups. However, gut-MMP9 levels were elevated in the MCAO-WT-proximal/distal groups with BT compared to the *naïve* and MCAO-WT-NBT groups, with no changes observed in the MCAO-TLR4-KO groups (Fig. 5f).

Taking together, these results suggest that infarct size may influence the risk of BT, which in turn increases infarct volume and GBD. Additionally, TLR4 deficiency reduces lesion size and prevents GBD and BT.

### Impact of TLR4 deficiency and infarct size in peripheral and central inflammation

The analysis of the effects of IS, BT, and TLR4 deficiency on peripheral inflammation revealed a reduction in T-CD4+ and CD8+ cells in the bone marrow of IS-TLR4-KO animals compared to the WT-proximal-NBT group, as well as an increase in blood monocytes in the WT-distal-BT group compared to the WT-proximal-NBT group (Fig. 6a, b). The evaluation of lymphocyte percentages in peripheral blood showed a trend toward lower levels in the WT-BT group compared to their respective NBT groups, with no differences observed between ischemic TLR4-KO animals and the WT groups (Fig. 6c). Inflammation in other organs showed a reduction in T-CD4+ and CD8+ cells in the spleen of the WT-distal-BT group compared to the WT-NBT group, and a decrease in all lymphocytes in the lungs of the TLR4-KO group compared to the other WT groups, both with and without BT (Fig. 6d, e).

**Figure 6.**
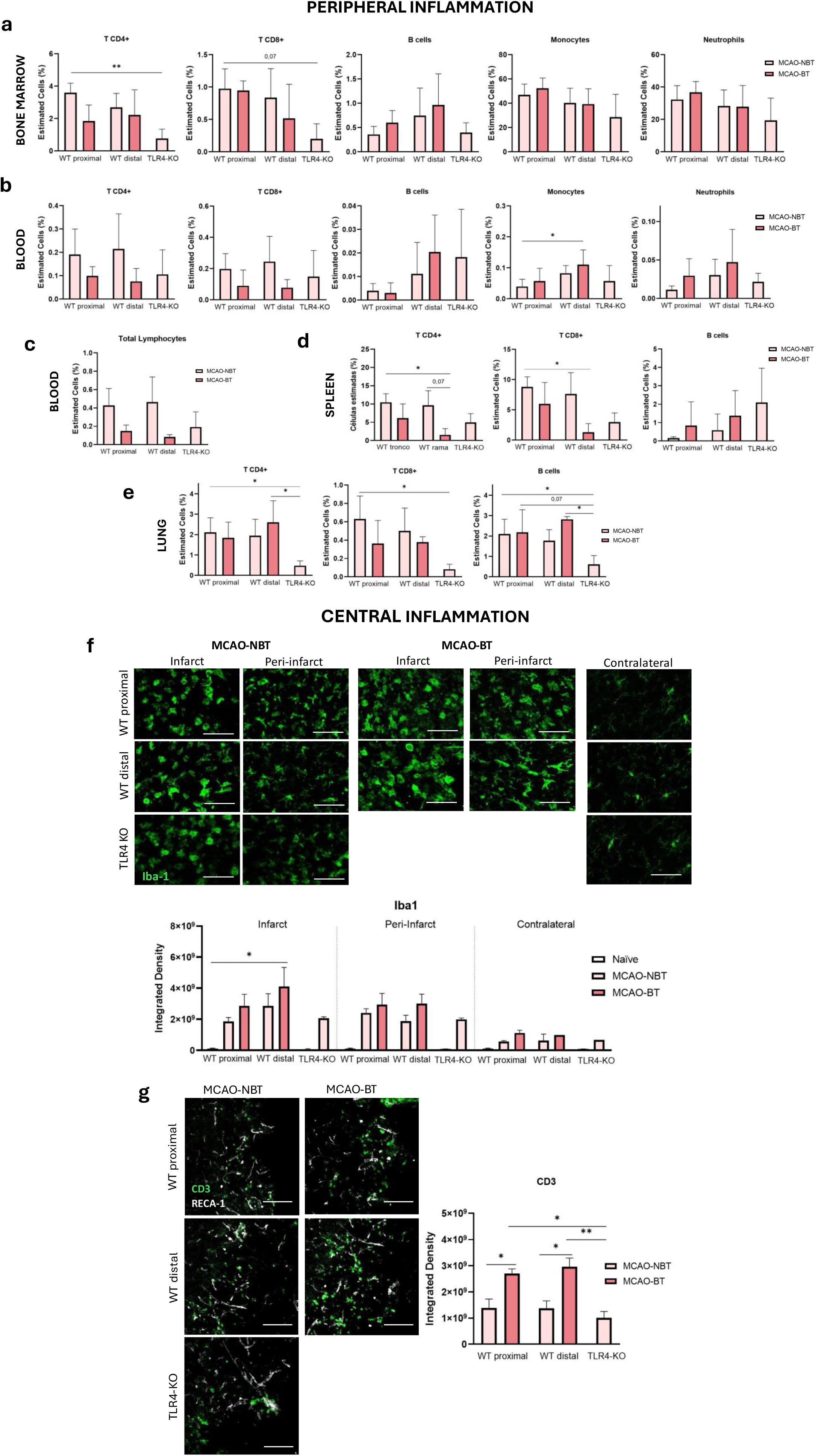
Peripheral and central immune response following IS in WT and TLR4-KO mice based on BT. **(a-e)** Immune cell levels in **(a)** BM, **(b,c)** peripheral blood, **(d)** spleen and **(e)** lung assessed by flow cytometry at 72h post-surgery in ischemic WT proximal (n (MCAO-NBT) = 9, n (MCAO-BT) = 4), WT distal (n (MCAO-NBT) = 8, n (MCAO-BT) = 4), and TLR4-KO (n=6) groups. Results are shown as mean ± SD; Kruskal-Wallis test followed by Dunn’s multiple comparisons test, \**p*<0.05, \*\**p*<0.01. **(f)** Microglial activation assessed by immunofluorescence at 72h post-stroke. **(f, top)** Representative confocal images of Iba-1 staining (green) in the infarct core, peri-infarct, and contralateral hemisphere in WT *naïve*, WT proximal MCAO-NBT and MCAO-BT, WT distal MCAO-NBT and MCAO-BT, and TLR4-KO naïve and ischemic groups (n=4-6). Scale bar: 50 μm. **(f, bottom)** Quantification of Iba-1 signal in the same regions expressed as mean ± SD; Kruskal-Wallis test followed by Dunn’s multiple comparisons test, * *p*<0.05. **(g)** T-cell infiltration in ischemic brain tissue assessed by immunofluorescence at 72h post-surgery. **(g, left)** Representative confocal images of the infarcted region in ischemic WT proximal MCAO-NBT and MCAO-BT, WT distal MCAO-NBT and MCAO-BT, and TLR4-KO (n=6). Staining for CD3 (green) and CD31 (white) are shown. Scale bar: 50 μm. **(g, right)** Quantification of T-cell infiltration in the lesion area expressed as integrated density. Results are shown as mean ± SD; Kruskal-Wallis test followed by Dunn’s multiple comparisons test, \**p*<0.05, \*\**p*<0.01.

Finally, the evaluation of central inflammation revealed an increase in microglial activation in the affected hemisphere of the WT-proximal-BT group compared to the *naïve* group, with no significant differences in the other groups. Additionally, T lymphocyte infiltration in the ischemic brain was significantly higher in the WT-BT group compared to their respective NBT groups and the TLR4-KO group (Fig. 6f, g).

## Discussion

One of the most common complications in stroke patients, affecting 30-60%, is the development of SAIs, such as stroke-associated pneumonia (SAP) and urinary tract infections (UTIs), which significantly worsen patient outcomes and increase mortality^13–15^. While dysfunctions and hospital management practices have been recognized as key factors in the development of these infections^16, 17^, recent experimental and clinical studies suggest that GBD and BT from the intestine to normally sterile organs, including the lungs, may also play a role in these complications^18–21^. However, despite these advances, the exact mechanisms underlying these processes and their impact on inflammation and brain damage remain insufficiently understood. In line with previous research, our experimental findings show at 72h after stroke, that a high percentage of animals exhibit both GBD and BT. Concerning GBD, we observed that animals with post-stroke BT had a more permeable and inflamed intestinal barrier, which was confirmed by histological analysis and, for the first time, through in vivo analysis of contrast agent permeability using MRI. These results confirm previous findings demonstrating the deleterious effect of stroke on the intestinal barrier, but our study introduces a novel non-invasive imaging technique to validate these observations.

Regarding BT, although typical intestinal bacteria were found in various abdominal organs, the most significant bacterial growth was observed in MLN. Previous studies suggest that post-stroke GBD facilitates bacterial translocation to the MLN, which acts as the first immune barrier between the intestine and the body, promoting the migration of bacteria to other organs via the lymphatic system^18, 31^. Detailed bacterial analysis revealed that the predominant phylum was *Bacillota*, while *Bacteroidota* was the least represented. This shift in the intestinal microbiota composition after stroke has been documented in earlier clinical and experimental studies^32–35^. Furthermore, 76% of animals with BT showed bacterial growth in the lungs, with *Enterococcus*, *Lactobacillus*, and *Escherichia coli* being the most prevalent bacteria, further confirming previous research showing the presence of typical intestinal bacteria in the lungs after stroke^19, 21^.

Concerning the inflammatory response 72h after the stroke, our data showed a reduction in T and B lymphocytes, along with alterations in monocyte and granulocyte levels in both the BM and blood of ischemic animals that developed BT. We also found a reduction in total lymphocyte levels in blood, indicative of lymphopenia in animals with BT. This phenomenon, already described in multiple experimental/clinical studies, would result from SI, which promotes the development of SAIs^36–38^. Our data on cytokines/chemokines in plasma revealed a significant increase in CINC-1, CCL20, and IL-10, along with a trend toward increased IL-1Ra in animals with stroke and BT. In particular, the findings on CINC-1 and CCL20, which are involved in the mobilization of monocytes and granulocytes during inflammation^39, 40^, would support the observed alterations in these cell populations in BM and blood in IS-BT animals. For IL-1Ra and IL-10, previous research has linked increased levels of these proteins to SI and increased susceptibility to developing SAIs^41–44^. Therefore, the higher levels of these molecules observed in MCAO-BT animals may indicate a greater vulnerability to post-stroke complications. Regarding central inflammation, our results revealed a significant increase in T-lymphocyte infiltration in the ischemic brain of animals with BT. This, coupled with the decrease in these cells in the blood of the MCAO-BT group, suggests that T-lymphocytes are being mobilized from the periphery to the ischemic brain. In this sense, previous studies have demonstrated that intestinal dysbiosis and GBD after stroke promote the activation and migration of T-lymphocytes to the ischemic brain, thereby intensifying brain inflammation^45–47^. Finally, the analysis of infarct volume indicated that alterations in peripheral and central inflammation associated with GBD/BT resulted in a significant increase in lesion size in animals with BT. These results are consistent with previous studies that found a higher incidence of infections in individuals with larger infarcts^13, 48, 49^. Additionally, we observed that the risk of BT increases as the infarct volume grows, suggesting that lesion size may also contribute to the development of these complications.

In various neurological diseases, such as autism, stress, and depression, sCD14 has been identified as a marker for GBD/BT processes^22–24^. In this context, we hypothesized that sCD14 could be a reliable marker for GBD/BT in stroke patients and its association with the development of infections. Our results indicated that larger infarct volumes, poorer collateral circulation, and worse neurological status correlated strongly with elevated sCD14 levels. Moreover, elevated sCD14 levels at 24h post-admission was strongly associated with an increased likelihood of developing in-hospital infections. In line with these findings, previous studies have shown that sCD14, along with lipopolysaccharide-binding protein (LBP), can moderately predict the development of SAP, particularly in older patients or those with worse neurological conditions at admission^25^. Furthermore, our results linking infarct volume and infection development are consistent with studies that show a direct relationship between the initial lesion size and the onset of complications^48, 49^. The analysis of sCD14 in our experimental model confirmed that animals with stroke and BT exhibited significantly higher sCD14 levels compared to other groups. Additionally, in this experimental model, we found a positive correlation between infarct volume and intestinal permeability to contrast agents, as determined by MRI, further supporting the hypothesis that GBD/BT processes may be occurring clinically, increasing the risk of SAIs, and that sCD14 could predict them early.

Given the pivotal role of the TLR4 in stroke, we investigated whether pharmacological inhibition using ApTOLL or its genetic absence could not only provide neuroprotection but also reduce GBD/BT following stroke. Our results showed that ApTOLL treatment significantly reduced infarct volume and peripheral inflammation, confirming its neuroprotective effects^6, 7, 50^. Regarding BT, pharmacological inhibition with a single dose of ApTOLL slightly reduced the process. However, when comparisons were made with a larger number of animals without treatment or with a vehicle, we observed a significant reduction in BT after stroke. Treatment with ApTOLL also led to changes in bacterial phyla, with the complete absence of the *Bacteroidota* and an increase in the *Actinobacterota* phyla. Furthermore, about the gut-barrier, we observed a reduction in MMP9 expression in the MCAO-ApTOLL-BT-group compared to the vehicle-treated-group. Although all these data suggest a beneficial effect of ApTOLL on GBD/BT, it would be necessary to increase the number of animals and/or doses to confirm this effect. Moreover, further research is needed to understand the impact of this single dose on the phyla translocated from the intestine. In terms of peripheral inflammation, we observed significant differences in the chemokines CX3CL1 and CXCL7 between treated-NBT and untreated-BT animals. The reduction of these chemokines, involved in leukocyte mobilization and intestinal damage during inflammation^51–54^, after ApTOLL treatment suggests a benefit of this drug not only at the peripheral level but also in the intestine. In the context of central inflammation, ApTOLL reduced granulocyte and T-lymphocyte infiltration in the ischemic brain compared to vehicle-treated groups, as previously described^50^.

Regarding the absence of TLR4 and its impact on GBD/BT, our results not only confirmed a reduction in infarct volume^3, 5^, but also showed a complete absence of BT in TLR4-deficient animals. Additionally, improvements in peripheral and central inflammation were observed in TLR4-deficient animals compared to WT animals, as previously described^4, 55, 56^, especially in those with BT. Concerning the gut, our data also showed a reduction of intestinal inflammation in TLR4-KO mice compared with WT mice. These findings suggest that TLR4 plays a crucial role in the processes of GBD/BT after stroke. Concerning the effect of the initial infarct size in WT mice on these processes, our data revealed no differences in the percentage of BT, intestinal barrier integrity, or peripheral and central inflammation.

In conclusion, this study demonstrated that stroke induces GBD and BT, leading to bacterial translocation to abdominal organs and even the lungs. These processes exacerbate both peripheral and central inflammation, thereby increasing infarct volume. Furthermore, we found that larger infarct volumes experimentally elevate the percentage of BT. Additionally, high levels of sCD14 in patients could not only indicate the presence of GBD following a stroke but also serve as an early predictor of the development of SAIs. Finally, we showed that pharmacological inhibition, and especially the genetic absence of TLR4, not only reduces infarct volume but also improves GBD and BT processes.

## Methods

Methods, supplementary results and any associated references are available in the online version of the paper.

## Acknowledgements

The authors would like to acknowledge David Segarra and M. Eugenia Zarabozo (AptaTargets S.L.) for their generous donation of the ApTOLL drug and vehicle to conduct this study.

This study has been supported by a grant from Spanish Ministry of Science and Innovation PID2020-117765RB-I00 (Dr Pradillo), from Leducq Trans-Atlantic Network of Excellence on Circadian Effects in Stroke TNE-21CVD04 (Drs Lo, Moro, and Lizasoain), from Instituto de Salud Carlos III and cofinanced by the European Development Regional Fund “A Way to Achieve Europe” PI23/00635, RICORS-ICTUS (Redes de Investigacion Cooperativa Orientadas a Resultados en Salud) RD21/0006/0001 and Programa FORTALECE-Instituto imas12, FORT23/00023 (Dr Lizasoain).

## Competing financial intestests

Macarena Hernandez-Jimenez is Scientific Advisor of the company AptaTargets S.L. The remaining authors declare no competing financial interests.

