## Supplementary material for "Impact of bacterial translocation in stroke outcome. Soluble CD14 as early clinical marker and effect of TLR4": Methods, supplementary tables 1-3 and figures 1-3

**ONLINE SUPPLEMETAL MATHERIAL**

### METHODS

#### Animals

All experiments were performed in male Wistar rats (RjHan:WI; Janvier Laboratories, France), male C57BL/6 (C57BL/6J Ola Hsd, WT mice, Envigo, Spain) and male TLR4-KO (B6(Cg)-*Tlr4*$^{tm1.2Karp}$/J Ola Hsd; Envigo, Spain) mice, 8 to 10 weeks old, and weighting 250-300g and 20-30g respectively. Animals were kept in ventilated cages at 22°C in a 12-hour light/dark cycle and 35% humidity with ad libitum access to food and water. All procedures were performed in accordance with the European Communities Council Directive (86/609/EEC) and approved by the Ethics Committee on Animal Welfare of University Complutense (PROEX No. 305/19 and 219.3/24) and are reported according to ARRIVE (Animal Research: Reporting of In Vivo Experiments) guidelines. A special effort was made to reduce the number of animals considering previous experience with these animals and using the statistical tool http://www.biomath.info. Mortality rate in our study was <7%.

#### Focal cerebral ischemia, drug administration and experimental groups

All the surgeries were conducted under anesthesia with isoflurane in a mix of $O_2$ and $N_2O$ (0.2/0.8 L/min). During surgery, body temperature was maintained at 37.0±0.5°C using a servo-controlled rectal probe–heating pad. In rats, a tandem permanent occlusion of the left common-carotid artery by ligature and the left middle-cerebral artery (MCAO) by electrocoagulation was performed as previously described[1, 2]. For the sham group, all the arteries were exposed but not occluded. To induce an IS in mice, only a permanent occlusion of the left MCA by electrocoagulation was performed as previously described[3, 4]. To study the effect of the lack of TLR4 and the lesion size in GBD/BT processes, in the TLR4-KO an in a group of WT-mice, the MCAO was done by electrocoagulation of the trunk of the artery just before its bifurcation between the frontal and parietal branches (WT-proximal occlusion). To obtain smaller infarct volumes, in another WT-mice group, the occlusion was performed in the same way, but more distally, occluding only the parietal branch of the MCA (WT-distal occlusion) as previously described[5].

The administration of vehicle or ApTOLL (Aptus Biotech, Madrid, Spain) in rats was performed 10min after the MCAO by intravenous injection using the tail vein and at a dose of 0.45mg/kg as previously described[6].

All groups were performed and quantified in a randomized fashion (coin toss) by investigators blinded to each specific treatment/condition. In rats, to determine the effect of IS on the GBD/BT development, the following experimental groups were used: 1) *Naïve* (n=8); 2) Sham (n=8) and 3) MCAO (n=28). To study the effect of ApTOLL in these processes after IS, we performed: 1) *Naïve*-Vehicle (n=4); 2) *Naïve*-ApTOLL (n=4); 3) MCAO-Vehicle (n=14) and 4) MCAO-ApTOLL (n=14). For mice, our experimental groups were: 1) *Naïve*-WT (n=4); 2) Naïve-TLR4-KO (n=4); 3) MCAO-WT-proximal (n=26); 4) MCAO-WT-distal (n=26) and 5) MCAO-TLR4-KO (n=10). Finally, after the analysis of bacterial growth in different peritoneal organs and lungs, all these experimental groups were divided into groups with BT or with no BT (NBT).

**Study Design and Patient Cohort**

This study included a total of 152 patients admitted to the stroke unit of Hospital 12 de Octubre between January 2020 and July 2024. All patients met the following general inclusion and exclusion criteria. Inclusion criteria: Patients aged over 18 years with an ischemic stroke, previously independent (pre-morbid modified Rankin Scale (mRS) < 2), and admitted within six hours of symptom onset or after a wake-up stroke. Exclusion criteria: Patients with transient ischemic attacks (TIA), lacunar strokes, intracranial bleeding secondary to trauma or subarachnoid hemorrhage, or with a history of stroke, acute myocardial infarction, severe systemic infection, or major surgery within the last three months were excluded. Additional exclusion criteria included active severe systemic inflammatory disease, pregnancy, or puerperium.

**Data Collection and Laboratory Variables**

Data on age, gender, cardiovascular risk factors (CVRF), prior medication, and baseline stroke severity (NIHSS) were collected for all patients. Laboratory data, including biochemistry, cell counts, and coagulation parameters, were obtained from admission blood samples. Additional clinical data, including NIHSS scores during hospitalization, infarct volumes, and 90-day mRS outcomes, were retrieved from discharge records and follow-up visits.

Infarct volumes were measured by neurologists using CT or MRI within the first 48 hours post-stroke and analyzed under double-blind conditions. Collateral circulation was evaluated using cerebral angiography (CTA or MRA) and graded with the Visual Collateral Score (vCS), ranging from 0 (absent) to 3 (good collaterals filling 100% of the occluded territory)[7].

**Blood Sample Collection and Biomarker Quantification**

EDTA-treated whole blood was collected 24h after admission. Platelet-poor plasma (PPP) was prepared by centrifugation to minimize platelet interference. Quantification of plasma zonulin and sCD14 was performed using commercially available enzyme-linked immunosorbent assay (ELISA) kits following the manufacturers' instructions: Human zonulin ELISA kit (Cusabio, ref: CSB-EQ027649HU), and Human sCD14 ELISA Kit (R&D Systems, ref: DC140).

**Statistical Analysis of the clinical data**

Baseline characteristics were stratified into tertiles according to sCD14 distribution (low, medium, high). Univariate analyses were performed to compare differences between tertiles. One-way ANOVA was used for continuous variables, assuming normal distribution, and results were reported as means ± SE. Categorical variables were compared using the Chi-square test.

Multivariable regression models were conducted to evaluate associations between sCD14 levels and clinical outcomes, including NIHSS, infarct volume, collateral circulation grade, and in-hospital infection. Linear regression was applied for continuous outcomes, and logistic regression for binary outcomes. Covariates included significant variables from univariate analysis (age, gender, dyslipidemia). Adjusted means and probabilities were calculated with 95% confidence intervals (CI) and visualized using bar plots or line graphs with error bars.

All analyses and visualizations were performed using R (v4.4.2), with ggplot2, dplyr, and broom as key packages. A p-value <0.05 was considered statistically significant.

**Sample Size Justification**

This is a pioneering study evaluating markers of intestinal barrier permeability in an Iberian stroke cohort. Due to the lack of prior research with these markers, a formal sample size calculation was not feasible. However, the study was designed to detect clinically meaningful differences with a power of 80% and $\alpha = 0.05$.

**Ethical Approval and Informed Consent**

This study was approved by the Institutional Review Board (IRB) of Hospital 12 de Octubre (CEIm: 20/604, following the latest revised version of the Helsinki Declaration from 2013). Informed consent was obtained from all patients or their legal representatives prior to inclusion in the study, in compliance with institutional and ethical guidelines for human research.

**Assessment of neurological deficits in our experimental model**

The neurological function was evaluated in *naïve*, and at 72h after surgery in MCAO-untreated and MCAO-Vehicle/ApTOLL rats by motor and behavioural scales[8]. For the motor test, animals were scored as: 0 points, no deficit; 1 point, failure to extend right forepaw fully; 2 points, decreased grip of right forelimb while tail pulled; 3 points, spontaneous circling or walking to contralateral side; 4 points, walks only when stimulated with depressed level of consciousness; 5 points, unresponsive to stimulation. For the behavioral characterization of the animals, first, assessment of each animal began with observation of undisturbed behavior in a clear plastic cage: body position (completely flat, 0 to upright position, 4) and spontaneous activity (none, 0 to repeated vigorous movement, 3). Then, animals were transferred to an arena for observation of the following behaviors: transfer arousal (coma, 0 to extremely excited, 5), gait (absolute incapacity, 0 to normal, 3), touch escape (none, 0 to extremely vigorous, 3) and positional passivity (no struggle when held with a hand, 0, maximal struggle, 4). For the motor test, the higher the score the worse the neurological function, while for the behavioral test the opposite occurs, the higher the score the better the outcome.

**Experimental infarct size and GBD determination by MRI**

In all animals subjected to experimental IS, infarct volume was assessed at 72h post-surgery using T2W-magnetic resonance imaging (MRI) (Icon 1T-MRI; Bruker BioSpin GmbH, Ettlingen, Germany). During the image acquisition, animals were anesthetized as previously described. T2WI3D images were acquired using a rapid acquisition with relaxation enhancement (RARE) technique, with the following parameters: repetition time (TR) = 2200 s, echo train length = 12, echo spacing = 15 ms (resulting in an effective echo time (TE) = 75 ms), number of averages = 1, field of view (FOV) = 34 × 31 × 12 mm. The acquired matrix size was 136 × 124 × 34 (resolution: 0.25 × 0.25 × 0.500 mm), and the total acquisition time was approximately 10 minutes. Infarct size, expressed as percentage of ischemic hemisphere (%IH) was determined as previously described[9].

The study of intestinal barrier permeability was performed in *naïve* and at 72h after IS in untreated rats using T1W abdominal MRI images enhanced by oral administration of a contrast agents. Imaging was conducted using a Bruker Biospec BMT 47/40 MRI system

(Bruker, Ettlingen, Germany) operating at 4.7T and equipped with a 12 cm gradient system. A 7 cm diameter radiofrequency probe was used for both transmission and reception. All animals underwent a 12-hour fast prior to imaging to optimize image quality and minimize intestinal peristaltic movements. During the imaging procedure, animals were anesthetized as previously described, and body temperature (maintained between 36.5 and 37.5°C) and respiratory rate were continuously monitored throughout the study. One hour prior to abdominal image acquisition, the animals received by oral gavage a solution containing 1.5 ml of mannitol (Manitol Mein 10%, Fresenius Kabi S.A.U ) and 0.5 ml of water as an osmotic agent, and a solution containing 250mg/kg of $MnCl_2·4H_2O$ (CAS number 13446-34-9, PanReac, Barcelona, Spain) dissolved in 1.5 ml of water as contrast agent[10]. Following this, 2D spin-echo T1-weighted images with fat signal suppression were acquired, synchronized with respiration to minimize motion artifacts from abdominal movement. The imaging parameters were as follows: TR = 802 ms, TE = 10 ms, number of accumulations = 8, field of view = 12 x 6 cm, acquisition matrix = 256 x 128, slice thickness = 1 mm, and 12 slices. This provided a planar resolution of 469 $\mu m^2$ and an estimated total scan duration of 13 minutes and 46 seconds. Finally, the acquired images were analyzed using ImageJ 1.41 software (NIH, Bethesda, MD, USA) to quantitatively assess damage to the intestinal barrier resulting from the leakage of the contrast agent into surrounding organs.

**Analysis of BT by microbiological cultures**

To investigate the development of BT after stroke, microbiological cultures of different peritoneal organs and lungs were performed in *naïve*, *naïve*+Vehicle, *naïve*+ApTOLL rats, as well as at 72h post-surgery in sham rats and all animal groups subjected to IS as described in the previous section.

Briefly, mesenteric lymph node (MLN), liver, spleen and lung samples were obtained aseptically from all animals under terminal anesthesia. Samples were weighed and mechanically homogenized until complete disintegration in cold sterile saline (1 ml for rat experiments and 0.5 ml for mouse experiments). Subsequently, a volume of 50 µl of the undiluted and diluted (1:10-100 in sterile saline) preceding solution was plated on Columbia agar (Becton, Dickinson and Company-BD, NJ, USA), MacConkey agar (BD), and Brucella agar (BD). The surgery and sample processing were performed under maximum sterile conditions in a biological safety cabinet.

Columbia agar and MacConkey agar plates were incubated at 35°C for 48 hours in standard atmospheric conditions for strict aerobic bacteria cultivation, while Brucella agar plates were incubated at 35°C for 96 hours in an anaerobic chamber to facilitate the growth of strict and facultative anaerobic bacteria. After incubation, bacterial colony-forming units (CFUs) were differentially counted by combining stereomicroscopy analysis and culturing and biochemical methods. Images of the cultures were captured and digitized using a stereomicroscope (SteREO Discovery V.8, with AxioCam ERc 5s Rev.2, Carl Zeiss Microimaging, Germany) and later processed using the Zen modular software (Carl Zeiss Microimaging). Colonies with different morphologies and opacity were marked from digitized fields and enumerated. The labelled colonies were picked and subcultured on Columbia or Brucella agar for 48 hours. Microorganisms were identified by biochemical tests (API systems, BioMerieux).
Results were expressed in $\log_{10}$CFU per gram of tissue. The detection limit was 1.3 $\log_{10}$ CFU/g (20 CFU/g).

The nomenclature used in the identification of the bacterial phyla found in our analysis of the BT after IS has followed the rules of the *International Code of Nomenclature of Prokaryotes* (ICNP)[11]. This new nomenclature is described in the following table.

| New nomenclature | Old nomenclature |
|---|---|
| *Bacillota* | *Firmicutes* |
| *Bacteriodota* | *Bacteroidetes* |
| *Actinomycetota* | *Actinobacteria* |
| *Pseudomonadota* | *Proteobacteria* |
| *Fusobacteriota* | *Fusobacteria* |

**Study of peripheral inflammation by flow cytometry and cytokine-array kit and levels of sCD14 by ELISA**

Peripheral inflammation was assessed in different *naïve* groups (see the corresponding figure legend) and at 72h post-experimental IS in blood, bone marrow (BM), MLN, spleen, and lung samples from both rats and mice using flow cytometry. In all samples, populations of CD4+ and CD8+ T lymphocytes, B cells, monocytes, and granulocytes/neutrophils were analyzed. Sample processing was performed similarly in rats and mice. Thus, with the exception of blood, all samples were mechanically homogenized in a 1% PBS-BSA solution (bovine serum albumin) and filtered using 50 µm cell strainers. After Fc blocking, red blood cell lysis, and the corresponding centrifugations, each sample was incubated with the antibody cocktail described in the following tables, using as negative controls the corresponding isotypic antibodies.

| Antibodies for Rat | | | |
|---|---|---|---|
| Cell population | Antibody | Fluorochrome | Supplier (Catalog No.) |
| CD4 T cells | CD45<br>CD4<br>CD3 | FITC<br>PE<br>PE | BioLegend (Cat. No. 202205)<br>BioLegend (Cat. No. 550002)<br>BioLegend (Cat. No. 201507) |
| CD8 T cells | CD45<br>CD3<br>CD8a | FITC<br>PE<br>APC | BioLegend (Cat. No. 202205)<br>BioLegend (Cat. No. 201507)<br>BioLegend (Cat. No. 201705) |
| B cells | CD45<br>CD24 | FITC<br>PE | BioLegend (Cat. No. 202205)<br>BD Biosciences (Cat. No. 562104) |
| Monocytes | CD45<br>CD11b/c | FITC<br>APC | BioLegend (Cat. No. 202205)<br>BioLegend (Cat. No. 201809) |
| Granulocytes | Anti-rat granulocytes<br>MHC Class II (I-Ek) | PE<br>APC | BD Biosciences (Cat. No. 554905)<br>Miltenyi Biotec (Cat. No. 130-107-874) |

| Antibodies for mice | | | |
|---|---|---|---|
| Cell population | Antibody | Fluorochrome | Supplier (Catalog No.) |
| CD4 and CD8 T cells | CD45<br>CD3<br>CD8<br>CD4 | FITC<br>Perp-Cy5<br>APC<br>PE | BD Biosciences (Cat. No. 553080)<br>BioLegend (Cat. No. 100218)<br>BioLegend (Cat. No. 100712)<br>BD Biosciences (Cat. No. 557308) |
| B cells | CD45<br>CD3<br>CD19 | FITC<br>Perp-Cy5<br>APC-Cy7 | BD Biosciences (Cat. No. 553080)<br>BioLegend (Cat. No. 100218)<br>BioLegend (Cat. No. 115530) |
| Monocytes and neutrophils | CD45<br>CD11b<br>Ly6G<br>Ly6C | FITC<br>APC<br>PE<br>PE-Cy7 | BD Biosciences (Cat. No. 553080)<br>BD Biosciences (Cat. No. 553312)<br>BD Biosciences (Cat. No. 551461)<br>BD Biosciences (Cat. No. 560593) |

After washing the stained cells, they were resuspended in 200 µl of FACS Flow (BD Pharmingen, USA) and 200,000 events were acquired per sample using a FACSCalibur flow cytometer with CellQuest software (BD Pharmingen, USA). The data of each cell population were obtained using FlowJoVR software (FlowJo, USA) and expressed as estimated cell percentage.

The study of pro- and anti-inflammatory cytokines and chemokines was conducted in rat plasma from *naïve* and at 72h in the MCAO-NBT and MCAO-BT rat groups using a cytokine array kit (R&D Systems Rat Cytokine Array, ref: ARY008). This sandwich-type immunoassay allows the simultaneous detection of 29 cytokines and chemokines. The protocol followed was provided by the company.

Finally, sCD14 levels as a marker of leaky gut were determined in plasma samples from *naïve* rats and at 72h post-surgery in the MCAO-NBT and MCAO-BT groups by ELISA (Cusabio, ref: CSB-E11178r). The protocol followed for its determination was that specified by the manufacturer.

**Analysis of central inflammation and the state of the intestinal barrier by immunofluorescence**

To assess central inflammation and intestinal barrier integrity/inflammation, rats and mice from *naïve* and at 72h post-surgery from experimental IS groups (see the corresponding legend for each figure) were transcardially perfused. Subsequently, brains and colon samples were collected, fixed, cut (30 µm of thickness), and stained following a previously described immunofluorescence protocol[12]. Central inflammation was evaluated on brain sections using free-floating immunofluorescence. Rabbit anti-Iba-1 (1:100, Palex ref. 019-19741) was used to analyze microglial activation, mouse anti-rat granulocytes (1:50, BD Biosciences ref. 554905) was used to assess granulocyte infiltration, and rabbit anti-CD3 (1:200, Abcam ref. ab5690) was used to evaluate T lymphocyte infiltration in the ischemic brain. Additionally, mouse anti-rat-RECA-1 (1:500, Biorad ref. MCA970R) and rat anti-mouse-CD31 (1:100, BD Biosciences ref. 553370) were used to label blood vessels. Antigens were visualized using the corresponding secondary antibodies labeled with Alexa Fluor 488 and Cy3. For cellular nuclei visualization, the TO-PRO-3 marker (1:10,000, Thermo Fisher Scientific, ref: S33025) was included in some stainings. Four random immunofluorescence images from the infarct or peri-infarct regions in five consecutive sections (separated by 720 μm) were acquired to cover the central and most affected part of the cerebral cortex in each animal, starting at +1.70 mm to -0.40 mm from bregma. Intestinal barrier integrity and inflammation were analyzed using on-slice immunofluorescence. Colon sections were incubated with primary antibodies rabbit-anti-ZO-1 (1:50, Invitrogen ref. 617300) and mouse-anti-MMP9 (Santa Cruz ref. Sc-393859). The corresponding secondary antibodies labeled with Alexa Fluor 488 and Cy3 were used for detection, along with TO-PRO-3 for nuclear staining. Immunofluorescence images were acquired similarly, with four random images from five different sections for each animal. All images were acquired as Z-stacks at 20x magnification via laser-scanning confocal microscopy (LSM710; Zeiss, Germany), and quantification was performed through densitometric analysis using ImageJ software.

**Statistical analysis of the experimental data**

Experimental results are presented as mean±standard deviation (SD). For the statistical analysis of the results, GraphPad Prism 8.0.1 software was used. Before starting the analysis, all our data were tested with Kolmogorov-Smirnov normality test, to study whether they fit to gaussian distributions. For parametric data, the Student-t test was used for comparisons between two groups, one-way ANOVA followed by Tukey's post-hoc multiple comparisons test, and two-way ANOVA followed by Bonferroni post-hoc tests for comparisons among more than two groups. Non-parametric data were analyzed using the Mann-Whitney U test for comparisons between two groups and the Kruskal-Wallis test followed by Dunn's post-hoc test for comparisons among more than two groups. The relationship between the infarct volume and the $MnCl_2$ post-contrast T1W MRI abdominal signal was evaluated using simple linear regression. Finally, categorical data were analyzed using the Chi-square test.

**SUPPLEMENTARY TABLE 1:** BT in naïve and at 72h in sham and MCAO rats

**a**

| ANIMAL | MLN | SPLEEN | LIVER | LUNG |
|---|---|---|---|---|
| Naïve1 | - | - | - | - |
| Naïve2 | - | - | - | - |
| Naïve3 | - | - | - | - |
| Naïve4 | - | - | - | - |
| Naïve5 | - | - | - | - |
| Naïve6 | - | - | - | - |
| Naïve7 | - | - | - | - |
| Naïve8 | - | - | - | - |
| Sham1 | - | - | - | - |
| Sham2 | - | - | - | - |
| Sham3 | - | - | - | - |
| Sham4 | - | - | - | - |
| Sham5 | - | - | - | - |
| Sham6 | - | - | - | - |
| Sham7 | - | - | - | - |
| Sham8 | - | - | - | - |

**b**

| ANIMAL | MLN | SPLEEN | LIVER | LUNG |
|---|---|---|---|---|
| MCAO1-BT | X | X | - | X |
| MCAO2-NBT | - | - | - | - |
| MCAO3-BT | X | - | X | X |
| MCAO4-NBT | - | - | - | - |
| MCAO5-BT | X | - | - | - |
| MCAO6-BT | X | - | - | - |
| MCAO7-NBT | - | - | - | |
| MCAO8-BT | X | - | - | X |
| MCAO9-BT | X | X | X | X |
| MCAO10-BT | X | X | X | X |
| MCAO11-BT | X | X | X | X |
| MCAO12-BT | X | X | X | X |
| MCAO13-BT | X | X | X | X |
| MCAO14-BT | X | - | X | X |
| MCAO15-BT | X | - | - | - |
| MCAO16-BT | X | - | X | X |
| MCAO17-BT | - | X | - | X |
| MCAO18-BT | - | X | - | X |
| MCAO19-NBT | - | - | - | - |
| MCAO20-BT | X | X | X | X |
| MCAO21-NBT | - | - | - | - |
| MCAO22-BT | X | - | - | X |
| MCAO23-BT | X | X | X | X |
| MCAO24-BT | X | X | X | X |
| MACO25-BT | X | - | - | - |
| MCAO26-NBT | - | - | - | - |
| MCAO27-NBT | - | - | - | - |
| MCAO28-NBT | - | - | X | - |

**Supplementary Table 2. Baseline Characteristics of Ischemic Stroke Patients Stratified by CD14 Circulating Levels at 24 Hours.** Baseline clinical and demographic characteristics of IS patients, categorized by sCD14 circulating levels (low, medium, high) at 24h post-admission. The table includes variables such as prior medical history, admission data, and treatment-related information. P-values were calculated to assess differences across sCD14 levels. Statistically significant differences ($p < 0.05$) are indicated in bold. CD14 levels were stratified into tertiles (33% per group).

| | sCD14 CIRCULATING LEVELS AT 24h ADMISSION | | | |
|---|---|---|---|---|
| | **low** | **medium** | **high** | **p-value** |
| mean value (SD) | 1288.81 (173.65) | 1802.39 (182.08) | 2511.04 (326.65) | |
| n | 51 | 51 | 50 | |
| **Age, mean (SD)** | 67.45 (15.73) | 72.86 (14.06) | 76.00 (14.11) | 0.014 |
| **Gender (Female), n (%)** | 19 ( 37.3) | 27 ( 52.9) | 32 ( 64.0) | 0.026 |
| **Previous mRS, n (%)** | | | | 0.383 |
| **mRS 0** | 42 ( 82.4) | 37 ( 72.5) | 38 ( 76.0) | |
| **mRS 1** | 7 ( 13.7) | 7 ( 13.7) | 9 ( 18.0) | |
| **mRS 2** | 2 ( 3.9) | 7 ( 13.7) | 3 ( 6.0) | |
| **Tobacco, n (%)** | | | | 0.326 |
| **Non smoker** | 30 ( 60.0) | 35 ( 70.0) | 39 ( 79.6) | |
| **Active smoker** | 13 ( 26.0) | 9 ( 18.0) | 6 ( 12.2) | |
| **Past smoker** | 7 ( 14.0) | 6 ( 12.0) | 4 ( 8.2) | |
| **Alcohol, n (%)** | | | | 0.162 |
| **Non Alcohol Use** | 38 ( 77.6) | 43 ( 84.3) | 44 ( 91.7) | |
| **Alcohol Use** | 8 ( 16.3) | 8 ( 15.7) | 3 ( 6.2) | |
| **Past Alcohol Use** | 3 ( 6.1) | 0 ( 0.0) | 1 ( 2.1) | |
| **Hypertension, n (%)** | 29 ( 56.9) | 38 ( 74.5) | 35 ( 70.0) | 0.144 |
| **Dyslipidemia, n (%)** | 23 ( 45.1) | 29 ( 56.9) | 35 ( 70.0) | **0.041** |
| **Diabetes, n (%)** | 18 ( 35.3) | 15 ( 29.4) | 12 ( 24.0) | 0.461 |
| **Atrial Fibrillation, n (%)** | 14 ( 27.5) | 9 ( 17.6) | 16 ( 32.0) | 0.240 |
| **Metallic Valve, n (%)** | 1 ( 2.0) | 0 ( 0.0) | 1 ( 2.0) | 0.599 |
| **Previous Ischemic Stroke, n (%)** | 7 ( 13.7) | 7 ( 13.7) | 3 ( 6.0) | 0.365 |
| **Previous Hematoma, n (%)** | 1 ( 2.0) | 0 ( 0.0) | 1 ( 2.0) | 0.599 |
| **Previous Myocardial Infarction, n (%)** | 7 ( 13.7) | 3 ( 5.9) | 2 ( 4.0) | 0.156 |
| **Arteriopathy, n (%)** | 4 ( 7.8) | 1 ( 2.0) | 2 ( 4.0) | 0.355 |
| **Chronic Kidney Disease, n (%)** | 5 ( 9.8) | 5 ( 9.8) | 0 ( 0.0) | **0.073** |
| **Previous Antihypertensive Therapy, n (%)** | 27 ( 52.9) | 37 ( 72.5) | 27 ( 54.0) | **0.076** |
| **Previous Lipid-Lowering Therapy, n (%)** | 23 ( 45.1) | 26 ( 51.0) | 27 ( 54.0) | 0.660 |
| **Previous Antiplatelet Therapy, n (%)** | 11 ( 21.6) | 13 ( 25.5) | 11 ( 22.0) | 0.876 |
| **Previous Anticoagulant Therapy, n (%)** | 13 ( 25.5) | 7 ( 13.7) | 13 ( 26.0) | 0.237 |
| **Admission Systolic Blood Pressure, mean (SD)** | 152.47 (22.33) | 153.09 (26.55) | 147.40 (26.29) | 0.505 |
| **Admission Diastolic Blood Pressure, mean (SD)** | 82.43 (14.61) | 80.78 (13.74) | 83.79 (14.00) | 0.591 |
| **Admission Creatinine, mean (SD)** | 0.94 (0.30) | 0.93 (0.36) | 0.94 (0.28) | 0.966 |
| **Admission Blood Glucose, mean (SD)** | 125.28 (42.78) | 140.83 (49.59) | 138.27 (55.40) | 0.252 |
| **Admission C-reactive Protein, mean (SD)** | 0.37 (0.50) | 0.72 (0.97) | 1.25 (2.70) | 0.057 |
| **Admission INR, mean (SD)** | 1.21 (0.56) | 1.04 (0.15) | 1.14 (0.32) | 0.085 |
| **Admission NIHSS, mean (SD)** | 10.20 (7.44) | 12.86 (8.06) | 15.70 (7.30) | **0.002** |
| **ASPECTS, mean (SD)** | 8.94 (1.13) | 9.06 (1.19) | 8.49 (1.45) | 0.133 |
| **Infarct Volume (cc), mean (SD)** | 18.69 (50.32) | 21.08 (44.06) | 62.30 (113.30) | **0.007** |
| **Hyperdense Artery, mean (SD)** | 7 ( 20.0) | 5 ( 17.9) | 6 ( 42.9) | 0.160 |
| **Laterality, n (%)** | | | | 0.988 |

| | | | | |
|---|---|---|---|---|
| Left side | 25 ( 52.1) | 26 ( 51.0) | 22 ( 45.8) | |
| Right side | 19 ( 39.6) | 22 ( 43.1) | 23 ( 47.9) | |
| Vertebrobasilar | 3 ( 6.2) | 2 ( 3.9) | 2 ( 4.2) | |
| Multiple | 1 ( 2.1) | 1 ( 2.0) | 1 ( 2.1) | |
| **Intra-arterial Occlusion, n (%)** | | | | 0.625 |
| MCA.M1 | 11 ( 26.8) | 13 ( 28.9) | 16 ( 35.6) | |
| MCA.M2 | 8 ( 19.5) | 12 ( 26.7) | 7 ( 15.6) | |
| Carotid T | 7 ( 17.1) | 8 ( 17.8) | 13 ( 28.9) | |
| Basilar | 1 ( 2.4) | 0 ( 0.0) | 1 ( 2.2) | |
| Vertebral | 0 ( 0.0) | 1 ( 2.2) | 0 ( 0.0) | |
| PCA | 3 ( 7.3) | 1 ( 2.2) | 1 ( 2.2) | |
| ACA | 1 ( 2.4) | 0 ( 0.0) | 0 ( 0.0) | |
| Others more | 10 ( 24.4) | 10 ( 22.2) | 7 ( 15.6) | |
| **Tandem, mean (SD)** | 6 ( 12.8) | 7 ( 15.6) | 5 ( 10.9) | 0.800 |
| **Collateral Circulation grade, mean (SD)** | 2.44 (0.64) | 2.44 (0.75) | 1.95 (0.81) | **0.007** |
| **Etiology** | | | | 0.392 |
| Atherothrombotic | 17 ( 33.3) | 19 ( 38.0) | 10 ( 20.0) | |
| Cardioembolic | 18 ( 35.3) | 14 ( 28.0) | 22 ( 44.0) | |
| Crytogenic | 6 ( 11.8) | 1 ( 2.0) | 4 ( 8.0) | |
| Unknown Cause | 5 ( 9.8) | 7 ( 14.0) | 7 ( 14.0) | |
| Incomplete Study | 2 ( 3.9) | 6 ( 12.0) | 4 ( 8.0) | |
| Unusual Cause | 3 ( 5.9) | 3 ( 6.0) | 3 ( 6.0) | |
| **tPA, n (%)** | 19 ( 39.6) | 20 ( 40.0) | 15 ( 31.2) | 0.603 |
| **Thrombectomy, n (%)** | 26 ( 53.1) | 35 ( 68.6) | 41 ( 85.4) | **0.003** |
| **Incomplete Reperfusion (< mTICI 3)** | 8 ( 33.3) | 14 ( 40.0) | 12 ( 33.3) | 0.808 |
| **Admission Leukocytes, mean (SD)** | 8.13 (2.27) | 9.18 (2.56) | 9.24 (3.30) | 0.079 |
| **Admission Neutrophils, mean (SD)** | 5.51 (2.38) | 6.51 (2.59) | 6.66 (3.31) | 0.084 |
| **Admission Lymphocytes, mean (SD)** | 1.81 (0.90) | 1.86 (0.70) | 1.72 (0.82) | 0.692 |
| **Admission Monocytes, mean (SD)** | 0.64 (0.20) | 0.65 (0.22) | 0.65 (0.24) | 0.914 |
| **Admission Platelets, mean (SD)** | 211.98 (55.35) | 220.20 (83.78) | 223.25 (59.28) | 0.692 |
| **Stroke Unit Leukocytes, mean (SD)** | 8.27 (2.31) | 8.56 (2.27) | 8.63 (2.43) | 0.727 |
| **Stroke Unit Neutrophils, mean (SD)** | 5.90 (2.29) | 6.33 (1.99) | 6.53 (2.26) | 0.370 |
| **Stroke Unit Lymphocytes, mean (SD)** | 1.59 (0.74) | 1.46 (0.56) | 1.46 (1.12) | 0.703 |
| **Stroke Unit Monocytes, mean (SD)** | 0.62 (0.18) | 0.67 (0.20) | 0.70 (0.25) | 0.163 |
| **Stroke Unit Platelets, mean (SD)** | 194.25 (45.92) | 207.51 (75.72) | 190.74 (46.40) | 0.346 |
| **Post-hemorrhage, n (%)** | 2 ( 4.3) | 7 ( 15.2) | 10 ( 22.2) | **0.045** |
| **Post-stroke Infection, n (%)** | 6 ( 11.8) | 17 ( 33.3) | 31 ( 62.0) | **<0.001** |
| **Stroke Unit NIHSS, mean (SD)** | 5.30 (6.41) | 7.45 (7.35) | 10.06 (7.94) | **0.008** |
| mRS 3 months | | | | 0.140 |
| mRS 0 | 16 ( 36.4) | 10 ( 21.7) | 6 ( 12.8) | |
| mRS 1 | 8 ( 18.2) | 9 ( 19.6) | 10 ( 21.3) | |
| mRS 2 | 5 ( 11.4) | 8 ( 17.4) | 4 ( 8.5) | |
| mRS 3 | 6 ( 13.6) | 10 ( 21.7) | 8 ( 17.0) | |
| mRS 4 | 5 ( 11.4) | 5 ( 10.9) | 6 ( 12.8) | |
| mRS 5 | 0 ( 0.0) | 1 ( 2.2) | 5 ( 10.6) | |
| All-cause mortality | 4 ( 9.1) | 3 ( 6.5) | 8 ( 17.0) | |

**Supplementary Table 3. Multivariable Analysis of 24-hour sCD14 Levels Adjusted for Dyslipidemia, Age, and Gender.** This table presents the results of multivariable regression analyses, including both linear and logistic models, as appropriate for the dependent variables. The analysis assesses the associations between 24h sCD14 levels (low, medium, and high) and clinical outcomes, adjusting for confounders including dyslipidemia, age, and gender. Model estimates, standard errors, T/Z ratios, and corresponding *p*-values for pairwise contrasts between sCD14 levels are provided.

| Dependent Variable | Term | Estimate | Standard Error | P-value | sCD14 levels comparison | Estimate | Standard Error | T/Z Ratio | P-value |
|---|---|---|---|---|---|---|---|---|---|
| **Admission NIHSS** | (Intercept) | 4,402 | 3,154 | 0,165 | low - medium | -2,666 | 1,574 | -1,693 | 0,211 |
| | CD14 medium | 2,666 | 1,574 | **0,093** | low - high | -5,113 | 1,617 | -3,162 | **0,005** |
| | CD14 high | 5,113 | 1,617 | **0,002** | medium - high | -2,447 | 1,529 | -1,6 | 0,249 |
| | Age | 0,094 | 0,045 | **0,041** | | | | | |
| | Female | -0,317 | 1,304 | 0,808 | | | | | |
| | Dyslipidemia | -1,043 | 1,368 | 0,447 | | | | | |
| **Stroke Unit NIHSS** | (Intercept) | -1,8 | 3,027 | 0,553 | low - medium | -1,682 | 1,536 | -1,095 | 0,519 |
| | CD14 medium | 1,682 | 1,536 | 0,275 | low - high | -4,023 | 1,558 | -2,583 | **0,029** |
| | CD14 high | 4,023 | 1,558 | **0,011** | medium - high | -2,342 | 1,488 | -1,574 | 0,26 |
| | Age | 0,116 | 0,045 | **0,010** | | | | | |
| | Female | -0,71 | 1,262 | 0,575 | | | | | |
| | Dyslipidemia | -0,758 | 1,323 | 0,567 | | | | | |
| **Infarct Size (cc)** | (Intercept) | -11,594 | 30,322 | 0,703 | low - medium | -5,766 | 15,317 | -0,376 | 0,925 |
| | CD14 medium | 5,766 | 15,317 | 0,707 | low - high | -48,278 | 15,691 | -3,077 | **0,007** |
| | CD14 high | 48,278 | 15,691 | **0,003** | medium - high | -42,511 | 15,285 | -2,781 | **0,017** |
| | Age | 0,675 | 0,441 | 0,129 | | | | | |
| | Female | -32,926 | 12,879 | **0,012** | | | | | |
| | Dyslipidemia | -6,559 | 13,38 | 0,625 | | | | | |
| **Collateral Circulation grade** | (Intercept) | 1,991 | 0,391 | **0,000** | low - medium | 0,105 | 0,196 | 0,538 | 0,853 |
| | CD14 medium | -0,105 | 0,196 | 0,592 | low - high | 0,635 | 0,193 | 3,283 | **0,004** |
| | CD14 high | -0,635 | 0,193 | **0,001** | medium - high | 0,53 | 0,174 | 3,05 | **0,008** |
| | Age | 0,004 | 0,006 | 0,462 | | | | | |

| | | | | | | | | | |
|---|---|---|---|---|---|---|---|---|---|
| | Female | 0,172 | 0,155 | 0,271 | | | | | |
| | Dyslipidemia | 0,234 | 0,164 | 0,156 | | | | | |
| **Post-Stroke Infection** | (Intercept) | -3,739 | 1,093 | **0,001** | low - medium | -1,182 | 0,544 | -2,170 | 0,076 |
| | CD14 medium | 1,182 | 0,544 | **0,030** | low - high | -2,41 | 0,551 | -4,375 | **0,000** |
| | CD14 high | 2,41 | 0,551 | **0,000** | medium - high | -1,228 | 0,431 | -2,849 | **0,012** |
| | Age | 0,027 | 0,015 | **0,073** | | | | | |
| | Female | 0,102 | 0,409 | **0,803** | | | | | |
| | Dyslipidemia | -0,358 | 0,423 | **0,397** | | | | | |

**SUPPLEMENTARY FIGURE 1 | Effects of ischemic stroke and ApTOLL treatment on neurological deficits and infarct volume**

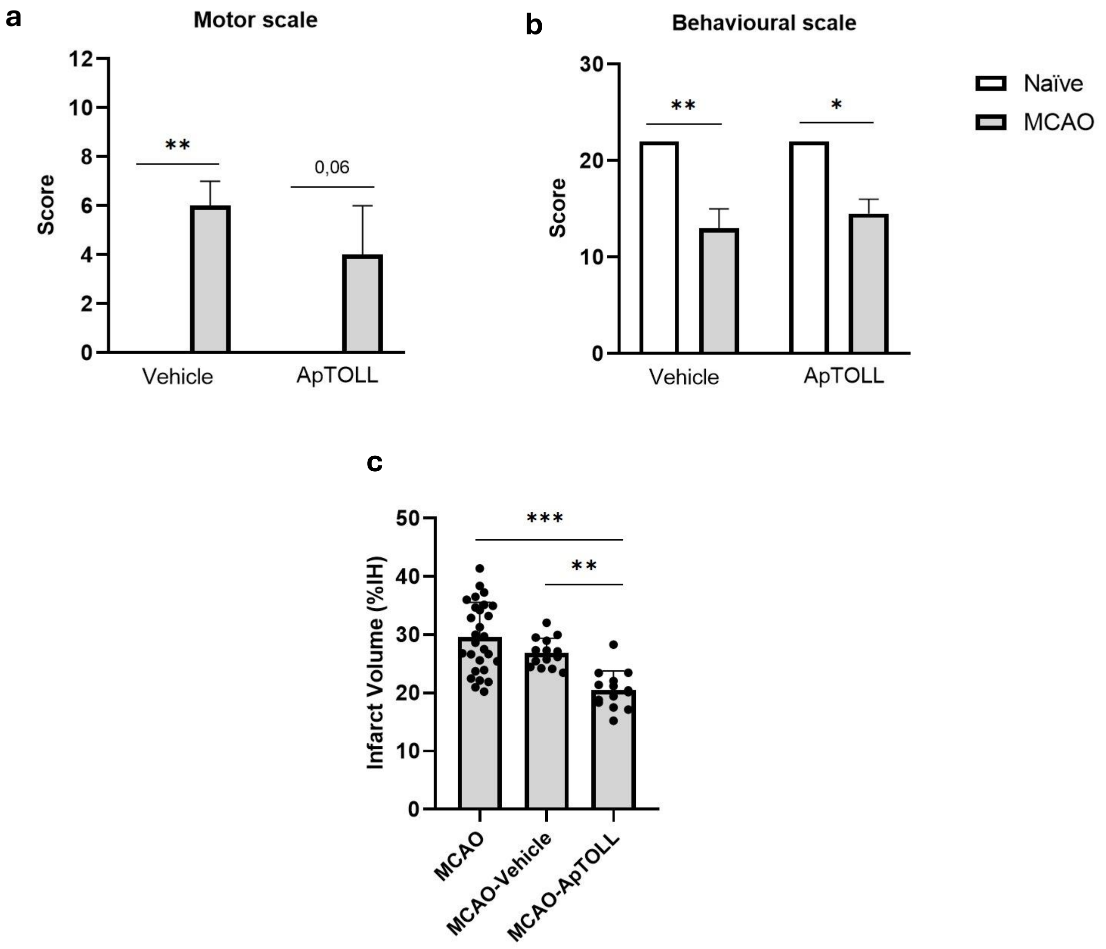


**Supplementary Figure 1: (a-b)** Neurological deficits were assessed in naïve and MCAO experimental groups treated with Vehicle or ApTOLL (n = 4 for naïve groups; n = 14 for MCAO groups). (a) Motor characterization and (b) behavioral characterization were evaluated. Results are expressed as Mean ± SD. Statistical analysis was performed using the Kruskal-Wallis test followed by Dunn's post hoc multiple comparisons test; *p < 0.05, **p < 0.01. **(c)** Infarct volume analysis at 72 h post-surgery in MCAO experimental groups from Block I (n = 28), comparing vehicle and ApTOLL treatments (n = 14 per treatment group). Infarct volume is expressed as the percentage of the infarcted hemisphere. Results are presented as Mean ± SD. Statistical analysis was performed using the Kruskal-Wallis test followed by Dunn's post hoc multiple comparisons test; **p < 0.01, ***p < 0.001.

### SUPPLEMENTARY FIGURE 2 | Influence of ApTOLL treatment and bacterial translocation on stroke outcome

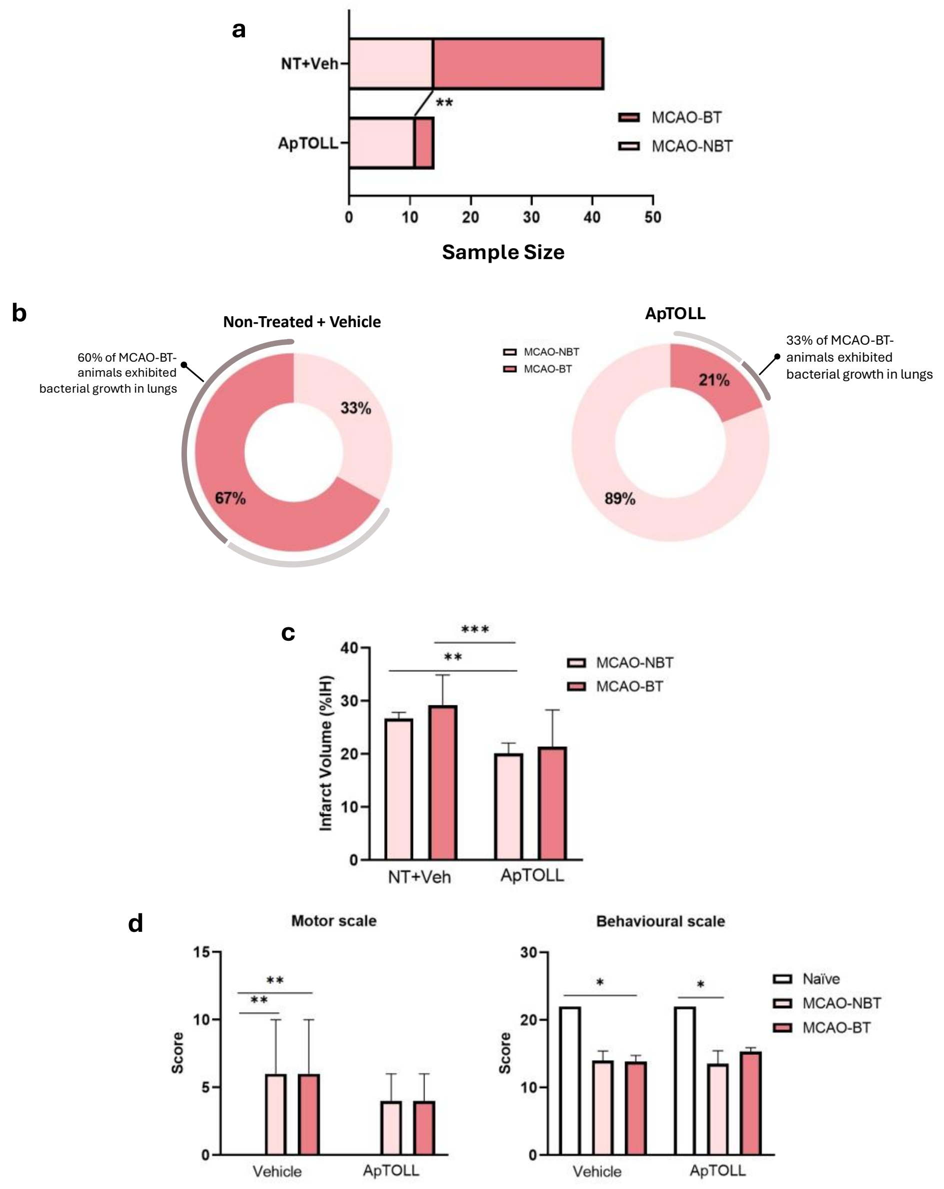


**Supplementary Figure 2: (a)** Proportion of ischemic animals according to treatment (Vehicle or ApTOLL) and bacterial translocation (BT) status at 72 h post-surgery. Groups compared: untreated + Vehicle ($n_{MCAO\text{-}NBT}$ = 14, $n_{MCAO\text{-}BT}$ = 28) and ApTOLL ($n_{MCAO\text{-}NBT}$ = 11, $n_{MCAO\text{-}BT}$ = 3). Statistical analysis performed using the Chi-squared test; **$p < 0.01$. **(b)** Percentage of ischemic animals developing (MCAO-BT) or not (MCAO-NBT) bacterial translocation at 72 h post-surgery in untreated + Vehicle ($n_{MCAO\text{-}NBT}$ = 14, $n_{MCAO\text{-}BT}$ = 28) and ApTOLL-treated ($n_{MCAO\text{-}NBT}$ = 11, $n_{MCAO\text{-}BT}$ = 3) groups. **(c-d)** Neurological deficits assessed in naïve animals treated with Vehicle or ApTOLL (n = 4 per group) and in ischemic animals stratified by bacterial translocation status (MCAO-NBT and MCAO-BT) and treatment (Vehicle: $n_{MCAO\text{-}NBT}$ = 6, $n_{MCAO\text{-}BT}$ = 6; ApTOLL: $n_{MCAO\text{-}NBT}$ = 7, $n_{MCAO\text{-}BT}$ = 3). (c) Motor characterization and (d) behavioral characterization were evaluated. Results are expressed as Mean ± SD. Statistical analysis was performed using the Kruskal-Wallis test followed by Dunn's post hoc multiple comparisons test; *$p < 0.05$, **$p < 0.01$.

**SUPPLEMENTARY FIGURE 3 | Immune cell levels in bone marrow and peripheral blood following ischemic stroke and ApTOLL treatment**

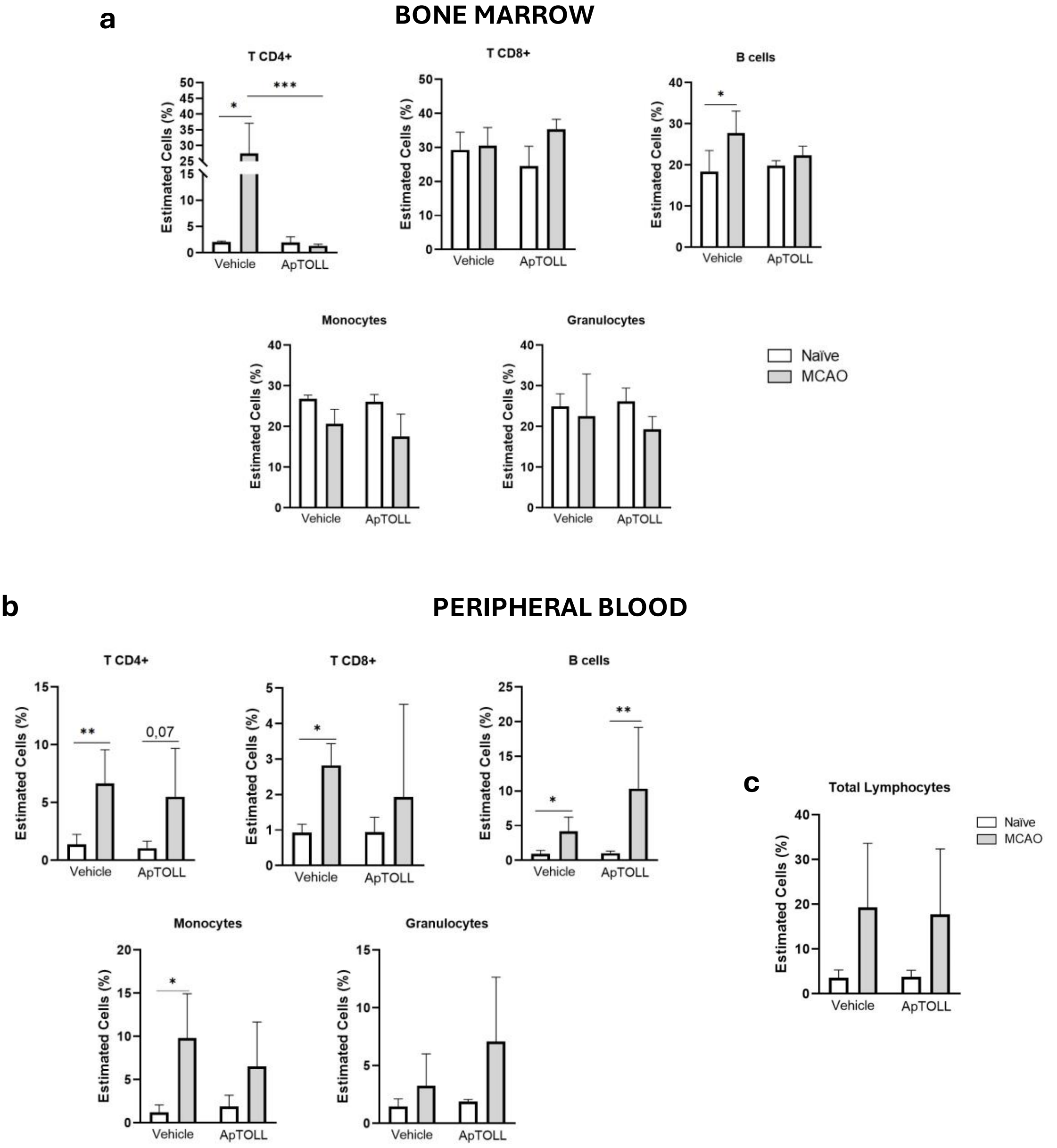


**Supplementary Figure 3: (a-c)** Immune cell levels in **(a)** bone marrow and **(b,c)** peripheral blood assessed by flow cytometry in naïve animals treated with Vehicle or ApTOLL (n = 4) and ischemic animals treated with Vehicle or ApTOLL (n = 14). Results are expressed as Mean ± SD. Statistical analysis was performed using the Kruskal-Wallis test followed by Dunn's post hoc multiple comparisons test; *p < 0.05, **p < 0.01, ***p < 0.001. Abbreviations**:** LB, B lymphocytes.
